# A CRISPR/Cas9-based enhancement of high-throughput single-cell transcriptomics

**DOI:** 10.1101/2022.09.06.506867

**Authors:** Amitabh C. Pandey, Jon Bezney, Dante DeAscanis, Ethan Kirsch, Farin Ahmed, Austin Crinklaw, Kumari Sonal Choudhary, Tony Mandala, Jeffrey Deason, Jasmin Hamdi, Azeem Siddique, Sridhar Ranganathan, Phillip Ordoukhanian, Keith Brown, Jon Armstrong, Steven Head, Eric J. Topol

## Abstract

Single-cell transcriptomics suffers from lapses in coverage of the full transcriptome, providing an incomplete gene expression profile of the cell. Here, we introduce single-cell CRISPRclean (scCLEAN), an *in vitro* molecular method that can be universally inserted into any single-cell RNA-seq workflow to improve the sensitivity of the assay. Utilizing CRISPR/Cas9, scCLEAN works to selectively remove highly abundant uninformative molecules, redistributing ~50% of reads to enrich for lowly expressed transcripts. Utilizing immune cells, we describe a validation of scCLEAN showing a 2.1-fold enrichment in library complexity with negligible off-target effects. Subsequently, applying scCLEAN to single-cell iso-seq samples results in a 4.6-fold improvement in unique isoform detection. Thus, demonstrating a benefit in short and long read sequencing applications. Finally, we illustrate the ability of scCLEAN to elucidate biological insights by applying it to two participant cohorts of cardiovascular samples, bringing to light novel molecular characteristics including inflammatory signatures.

## Introduction

Full transcriptome analysis has brought many molecular mechanisms to the forefront of translational investigation of disease processes. Similarly, gene expression profiling at single-cell resolution has revolutionized scientists’ ability to investigate biology, enabling the examination of inter- and intra-population heterogeneity^1^, developmental transition states^2^, regulatory mechanisms^3,4^, and cell-cell communication networks^5,6^. Due to recent technological breakthroughs, single-cell RNA sequencing (scRNAseq) experimental scope can now encompass millions of cells, allowing for the generation of detailed atlases of human disease^7–9^.

However, while massively parallel scRNAseq has improved the scale of data collection, similar gains have not yet materialized in analytical pathways. In turn, an increase in quantity of data has not necessarily translated to a concomitant increase in quality. Data generated via scRNAseq suffers from higher levels of noise due to technical variation and batch effects, input material quantity, amplification bias, read dropouts, and gene expression stochasticity^10–17^. Collectively, a detection efficiency from 3% to 25% exists^18–22^. As a result, only a small fraction of the nascent mRNA is processed, potentially confounding biological interpretation and reproducibility. While computational approaches are being developed to combat this problem inherent to scRNAseq^23–25^, an effective molecular solution remains to be elucidated.

As scRNAseq utilizes relatively low input, a high number of PCR cycles are required^26,27^, and less abundant molecules are often outcompeted by more abundant species during amplification^28–30^. Thus, the ideal application for CRISPR/Cas9 guided improvement in scRNAseq lies in steps prior to library preparation, allowing capture of less abundant molecules more frequently. To overcome these current bottlenecks in scRNAseq, we present a molecular solution, termed single-cell CRISPRClean (scCLEAN), that functions inversely to capture-based enrichment by removing unwanted sequences of high abundance. Since scCLEAN is applied to fully prepared cDNA, it is a simple turnkey solution that can be incorporated into any single-cell RNA-seq protocol, supporting widespread application.

Herein we demonstrate the utility and downstream analytical benefit of scCLEAN. We leveraged publicly available single-cell datasets to identify targets for depletion that could be applicable to most, if not all, source cell and tissue. After identification of candidate targets, we performed *in silico* modeling to validate the theoretical boost in sensitivity. Thereafter, we experimentally validated scCLEAN on peripheral blood mononuclear cells (PBMCs), a well characterized tissue with multiple distinct cell types^31^, demonstrating an improvement in cell type identification. Subsequently, to evaluate applicability to actionable transcriptome targets, scCLEAN was applied on PBMC derived single-cell iso-seq^32,33^. Finally, to highlight the ability of scCLEAN to enhance biological interpretation, we applied the approach to two separate cohorts of primary cells isolated from coronary and pulmonary artery locations: vascular smooth muscle cells (VSMCs) and vascular endothelial cells (VECs). In a notoriously difficult system to characterize due to the functional plasticity of smooth muscle cells^34^, scCLEAN enables a clear distinction between coronary and pulmonary arteries in addition to delineating key molecular differences in inflammatory signaling.

## Results

### Design and in silico Validation

To establish a framework of candidate transcripts that could safely be removed from a scRNAseq analysis without influencing downstream analysis, we focused on the 10x Genomics Chromium platform due to its widespread use^35^. A cohort of 14 publicly available scRNAseq datasets generated with Chromium v3.1 technology were benchmarked to examine consistent trends in read distribution across a spectrum of human samples (Supplementary Table 1). We found that, despite mRNA selection via polyA priming in the Chromium workflow, many resulting reads are consistently dropped from analysis because they do not align to the transcriptome (Fig. 1a). Thus, creating an opportunity to employ CRISPR/Cas9 depletion to evaluate the impact on single-cell data.

**Fig. 1:**
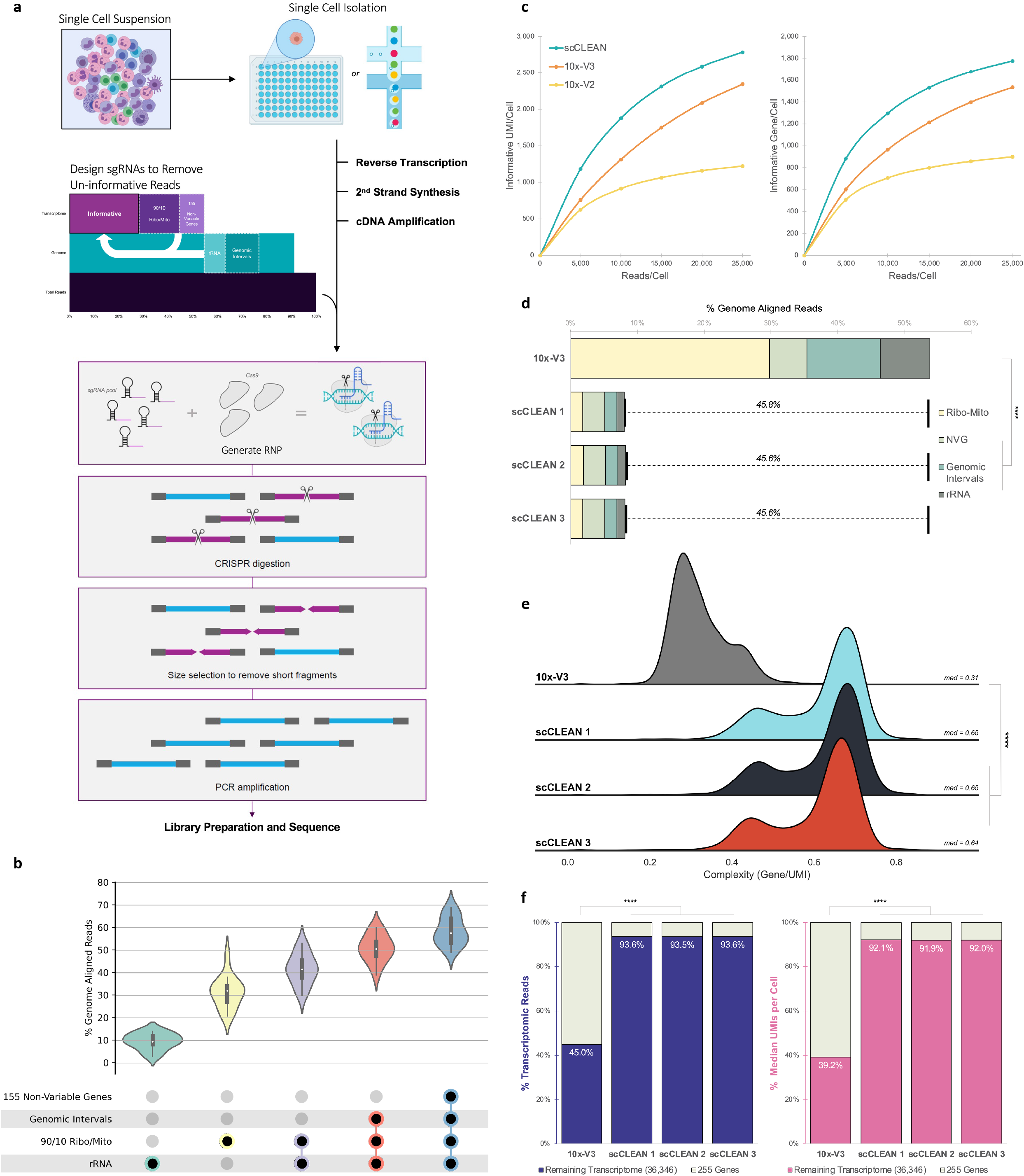
Evaluation of scCLEAN impact on single cell transcriptomic sensitivity. **a**, Schematic representing the insertion of CRISPR/Cas9 depletion into the standard single cell RNAseq (scRNAseq) workflow. A sgRNA pool is designed targeting 4 distinct regions: genomic intervals, non-polyadenylated rRNA (rRNA), 90 ribosomal and 10 mitochondrial genes (Ribo/Mito), and 155 non-variable housekeeping genes (NVG), redistributing reads to informative molecules. **b**, Distribution of genomic aligned reads attributed to each of the 4 targeted regions across 14 datasets. In total, 57% of aligned reads are targeted for redistribution. **c**, *In silico* modeling compares median genes per cell and median UMIs per cell as a function of sequencing depth. Uninformative molecules associated with the 255 targeted genes were ignored. **d**, Percent of genomic reads within PBMC samples (n=4) that aligned to each of the 4 targeted regions, on average depicting a read re-distribution of 46%. **e**, Distribution of cell complexity (gene/UMI), or the rate of gene recovery per unique molecule, for each of the 4 PBMC samples, quantifying a 2.1-fold boost in median complexity (med). **f**, Left, the ratio of transcriptome aligned reads per cell associated with the 255 targeted genes (tan) versus the remaining 36,346 genes in the transcriptome (blue), which increased from 45% (10x-V3) to 94% (scCLEAN). Right, the ratio of median UMIs per cell associated with the 255 targeted genes (tan) versus the remaining 36,346 genes in transcriptome (pink), which increased from 39% (10x-V3) to 92% (scCLEAN).

We constructed a single-guide RNA (sgRNA) library that targets two distinct categories of intervals: genomic and transcriptomic. Reads that align to the genome, but not the transcriptome, are ideal candidates for removal because they are computationally filtered before any downstream processing in the scRNAseq workflow. These targeted genomic regions consist of non-protein-coding rRNA and conserved genomic intervals, contributing 10% and 9% to the total aligned read count, respectively (Fig. 1b). In addition to genomic reads, 255 highly abundant protein-coding genes were also targeted for depletion given their expression varies little across different tissue types (see Supplementary Methods). Collectively, the 225 genes (90 protein-coding ribosomal genes, 10 protein-coding mitochondrial genes, and 155 protein-coding non-variable genes) constitute 39% of aligned reads (Fig 1b).

By removing uninformative molecules, we aimed to maximize read allocation to lower expressed genes, which tend to have greater biological relevance. As an initial validation and proof of concept of read redistribution, we performed *in silico* modeling (see Supplementary Methods). To properly compare the boost in per cell sensitivity, the reads and UMIs associated with the 255 targeted genes were ignored (hereinafter referred to as uninformative molecules), resulting in an average boost in both informative median UMIs/cell (35%) as well as genes/cell (29%) (Fig. 1c). Thus, confirming that the selective removal of highly abundant molecules, not contributing to downstream analyses, is in fact, a valuable methodology to enhance single-cell sensitivity.

### Experimental Validation in PBMCs

As a proof-of-principle demonstration, scCLEAN was experimentally validated on a cohort of primary PBMCs prepared with the standard Chromium 3’ v3.1 workflow (10x-V3) (n=1) and scCLEAN-treated samples (scCLEAN) (n=3), each comprised of roughly 11,000 cells. scCLEAN redistributed 46% of reads to other molecules in the single-cell library, representing an aggregated depletion percentage of 85% (Fig. 1d). Alignment to the standard 10x transcriptome reference illustrates a 2.1-fold increase in the complexity of the library (Fig. 1e). For every 1,000 UMIs detected per cell, 310 genes were identified in the 10x-V3 condition, whereas 650 genes were identified as a result of scCLEAN (Fig. 1e). Furthermore, scCLEAN reconfigured the composition of reads per cell. Instead of the non-targeted genes (36,346) accounting for a mere 39% of all UMIs per cell, scCLEAN greater than doubled (2.4x) that metric to 92% (Fig. 1f). In addition to a compositional change, scCLEAN statistically increased the total number of informative UMIs and genes identified per cell (Supplementary Fig. 1a). Collectively, scCLEAN achieves the same information as the standard Chromium workflow but with a 2-fold reduction in sequencing depth (Fig. 2a).

**Fig. 2:**
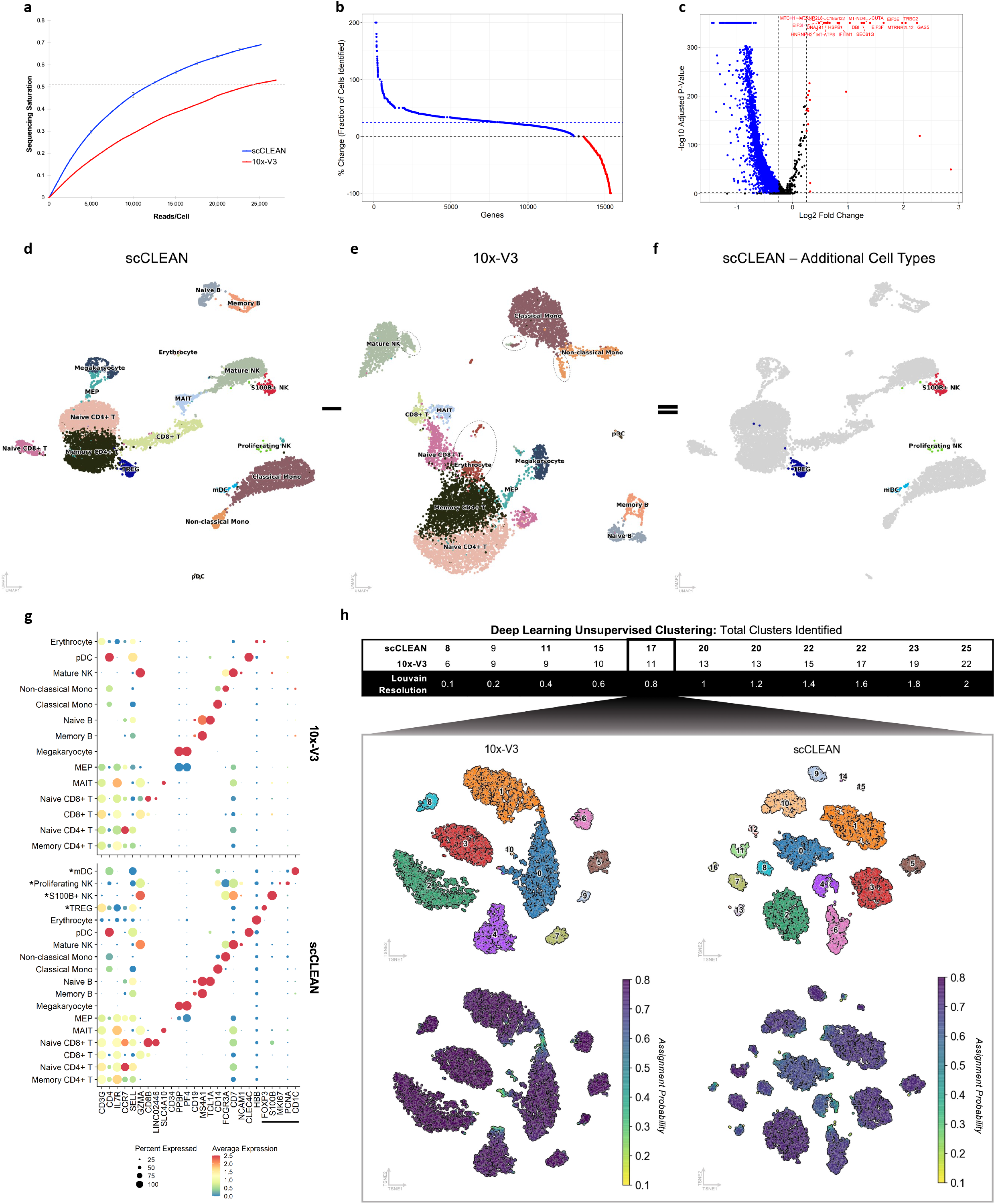
Enhanced clustering resolution contributing to robust cell type characterization. **a**, Sequencing saturation as a function of read depth comparing scCLEAN (blue) and 10x-V3 (red). Dotted line intersects the 10x-V3 condition at the gold standard sequencing depth of ~25,000 reads per cell, crossing the scCLEAN condition at roughly half. **b**, Illustrates the effect of scCLEAN (blue) on gene dropout relative to 10x-V3 (red). Of the genes detected in both conditions, the percent change in the total number of cells where each gene was identified. Blue dotted line represents the average boost in gene detection (24%). **c**, Volcano plot of differential expression analysis performed on all genes found in at least 10% of cells. Genes that passed a p-value of 0.05 and log2 fold change of 0.25 associated with scCLEAN (blue) (4,896) and 10x-V3 (red) (33). **d-f**, Cell clustering at the entropy (ROGUE) optimized resolution utilizing automatic cell type identification (CIPR) representing **d** scCLEAN, **e** 10x-V3 (dotted circles denote the location of cell types identified with scCLEAN but not characterized with 10x-V3), and **f** the clusters and cell types exclusively identified with scCLEAN (4 additional cell types). **g**, Expression of cell markers from a FACS sorted reference dataset for each of the PBMC cell types from each condition: 10x-V3 (top) and scCLEAN (bottom). Cell markers for the four additional cell types (*) annotated with a black line. **f**, Clustering output from an unsupervised deep learning algorithm (DESC) spanning the entire range of potential Louvain resolution inputs. Top, TSNE clustering plots from a chosen resolution (0.8) identifies 11 clusters with 10x-V3 and 17 clusters with scCLEAN. Bottom, paired assignment probability representing the confidence of assigning each cell to a specific cluster.

Next, we sought to elucidate and quantify read redistribution at the gene level. Of the 22,272 total genes detected, 19,703 (88%) exhibited an increase in median counts of genes/cell with scCLEAN (Supplementary Fig. 1c). Despite observation of a significant boost with scCLEAN, gene expression between scCLEAN and 10x-V3 remained in high concordance with a 0.95 coefficient of determination (Supplementary Fig. 1c). We subsequently performed differential expression analysis to gain a deeper insight into gene detection. Of the 6,101 genes expressed in at least 10% of cells, 4,896 genes (80%) were enriched in the scCLEAN condition and were expressed in ~24% more cells (Fig. 2b), consequently reducing gene dropout. No change was observed in 1,172 genes (19%), and 33 genes (1%) showed a significant enrichment in the 10x-V3 condition (Fig. 2c). While 33 genes passed the significance threshold, only 21 were determined to be off-target and 18 were direct targets of scCLEAN guides (Supplementary Fig. 1d-e).

Mitochondrial (MT) genes and the fraction of counts of MT genes per barcode is a standard quality control metric to identify dead or decaying cells in scRNAseq workflows^36,37^. However, analysis of the 14 datasets suggested that of these MT gene sets, 10 could be candidates for removal, and in turn reassignment of their reads in the scCLEAN protocol. We therefore aimed to verify that an MT-filtering based method could still confidently identify dead cells without substantial variation of QC metrics for scRNAseq analysis. A scRNAseq library representing PBMC cells consisting of a 30% dead cell population was generated. Subsequently, dead cell removal was performed on the sequencing data using standard and scCLEAN protocols, confirming the ability to filter the true dead cell population with up to 75% accuracy (Supplementary Fig. 2a-f).

For the achieved benefit observed with increased transcript detection, a similar improvement in biologically relevant signal is necessary, effectively resulting in a boost of biological signal versus noise. Using random matrix theory to parse between signal and noise within the data we observe that scCLEAN maintains sample variance (signal) while simultaneously reducing the false discovery rate (noise) as more genes are included in secondary analysis (Supplementary Fig. 3a-b). Consequently, by reducing noise, scCLEAN improves single-cell analysis by affecting a crucial step of data interpretation: dimensionality reduction (DR)^38,39^. Using a statistically principled optimization algorithm, evaluation of DR performance indicated that irrespective of the technique (PCA, diffusion, or autoencoder), scCLEAN retains greater biological variance within the latent representation (Supplementary Fig. 3c-d).

Our results indicate that scCLEAN increases transcriptomic sensitivity, reduces read dropouts, and boosts signal to noise. However, the end benefit must be an improvement in biologically relevant data obtained through scRNAseq. Therefore, we next assessed the performance of scCLEAN on cell clustering and subsequent cell-type identification. At the optimized clustering resolution (Supplementary Fig. 1b), the scCLEAN condition identified four additional cell clusters (Fig. 2d-f), due to the recovery of cell markers that had negligent expression within the 10x-V3 condition (Fig. 2g). These results were consistent across three replicates representing distinct samples of cells (Supplementary Fig. 1f). It is important to note that while the cells composing the four additional clusters existed within the 10x-V3 condition, given the resolution of analysis and decreased signal to noise ratio present, they simply couldn’t be transcriptionally distinguished as distinct clusters (Fig. 2e). Finally, using a fully unsupervised deep learning algorithm, to avoid any potential bias from the previous cell clustering, scCLEAN consistently identified more clusters facilitating additional characterization of heterogenous biology and rare cell type identification (Fig. 2h). To illustrate the improved characterization of cluster identity, genes were jointly embedded onto the latent space alongside cells. All four additional scCLEAN cell types had more closely associated genes in latent space according to Euclidean distance, revealing a potential explanation for improved characterization (Fig. 3a-b). For mucosal-associated invariant T (MAIT) cells, a cell type identified in both conditions, only 12 genes passed a cell type specificity threshold in the 10x-V3 condition whereas 575 genes passed the same threshold with scCLEAN (Supplementary Fig. 4a-e). In addition, cell markers for all cell types in the scCLEAN condition display a statistically significant higher cell type specificity according to four separate metrics (Gini index, max value, standard deviation, and entropy) (Supplementary Fig. 1g), illustrating the enhanced ability to distinguish between cell types and states with scCLEAN.

**Fig. 3:**
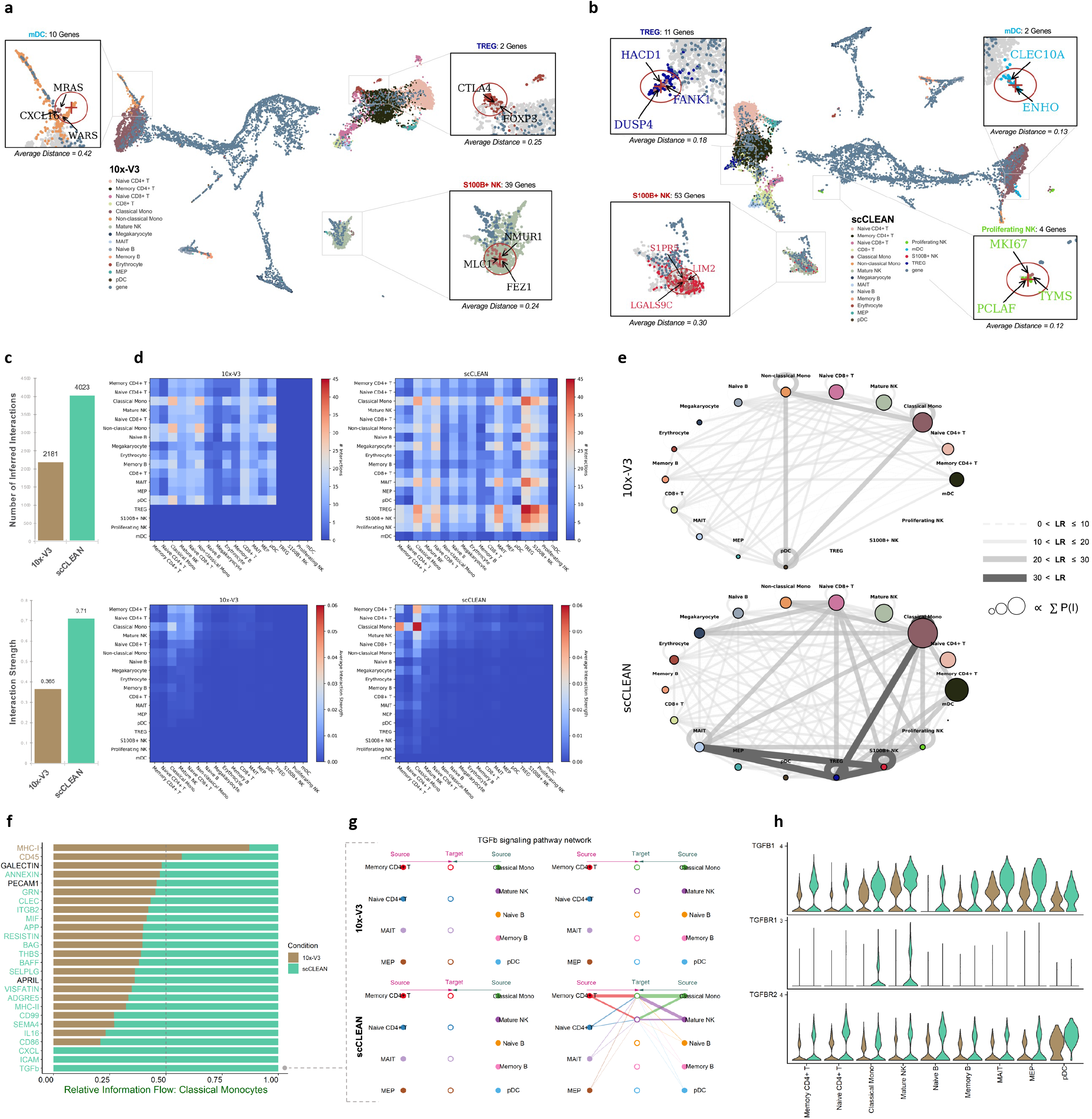
Increased cell type specificity improves cell to cell communication network clarity. **a**, Co-embedding of genes onto 10x-V3 latent space (SIMBA). Spatial proximity between genes and cell type quantifies specificity. Gene query with a radius of 0.5 UMAP coordinates from the spot of highest cell density according to where each cell type should be identified (spatial similarity with scCLEAN). The location of proliferating NK cells could not be determined. Average Euclidean distance to the center of each query. **b**, Co-embedding of genes onto scCLEAN latent space. Gene coloring represents a positive cell type identification. **c**, Top, total number of inferred cell to cell communication interactions between 10x-V3 (brown) (2,181 interactions) and scCLEAN (green) (4,023 interactions). Bottom, average interaction strength (1.9-fold increase). **d**, Top, signal heat matrix of total interactions with cell type specificity comparing 10x-V3 (left) with scCLEAN (right). Bottom, signal heat matrix of interaction strength. **e**, Network diagram depicting cellular interactions from 10x-V3 (top) and scCLEAN (bottom). Edge weights calculated according to the total number of ligand-receptor interactions associated with each cell type. Node size is proportional to the sum of all inferred probabilities between cell types, or the strength of an interaction occurring. **f**, Relative information flow between 10x-V3 (brown) and scCLEAN (green) of biological pathways associated with classical monocytes. Colored pathways reflect passing a Wilcoxon signed-rank test (p<0.05). 21/26 pathways statistically enhanced with scCLEAN. **g**, Network diagram depicting directional interactions of the TGFb signaling pathway between cell types of 10x-V3 (top) and scCLEAN (bottom). **h**, Gene expression between 10x-V3 (brown) and scCLEAN (green) of the main TGFb ligand (TGFB1) as well as both co-receptors (TGFBR1 and TGFBR2) necessary for an interaction to be inferred.

To highlight the biological benefit gained with scCLEAN, we further examined its role on the inference of cell-cell communication. scCLEAN increased the total number of ligand-receptor (LR) interactions by 1.8x and the average strength of interaction by 1.9x (Fig. 3c), enhancing the resolution of cellular communication networks (Fig. 3d-e). We next determined the impact of enhanced interaction resolution on comprehensive signaling pathways and focused on classical monocytes considering they had the strongest inference probability in both conditions (Fig. 3e) and play an integral role in inflammatory cytokine communication^40^. 21 out of the total 26 (80%) identified signaling pathways were statistically enriched with scCLEAN (Fig. 3f). Interestingly, three pathways were only identifiable as a result of scCLEAN: CXCL, ICAM, and TGFb. The TGFb pathway requires the primary TGFB1 ligand, as well as both TGFBR1+TGFBR2 coreceptors, to be expressed for an interaction to occur. In the 10x-V3 condition, there is insufficient expression levels of TGFBR2 (Fig. 3g-h). However, with scCLEAN, classical monocytes and natural killer cells have noticeable expression of TGFBR2 (Fig. 3g-h). A communication pathway was identified because a lowly expressed cofactor in a heteromeric complex was detected above the statistical threshold only with scCLEAN.

### Single-cell Iso-Seq

To investigate the broad applicability of the method, scCLEAN was next applied to PBMC derived single-cell iso-seq datasets. Despite similar quality control metrics (Supplementary Fig. 5a-c) and slightly fewer aligned reads in the scCLEAN condition (Fig. 4a), scCLEAN increased the total unique informative isoforms (not associated with the 255 targeted genes) from 17,131 (10x-V3) to 78,935 (scCLEAN). Novel isoform identification increased from 4,575 (10x-V3) to 19,112 (scCLEAN) correlating with a 4.6- and 4.2-fold improvement, respectively (Fig. 4b). More importantly, as sequencing depth increased, unique isoform identification plateaued in the 10x-V3 condition (slope=0.02) but continued to scale linearly with scCLEAN (slope=0.15), fundamentally altering the rate at which unique isoforms are detected (Fig. 4c). Given substantial increase in isoform detection, we next assessed whether the increase in throughput with scCLEAN also improves single-cell analysis.

**Fig. 4:**
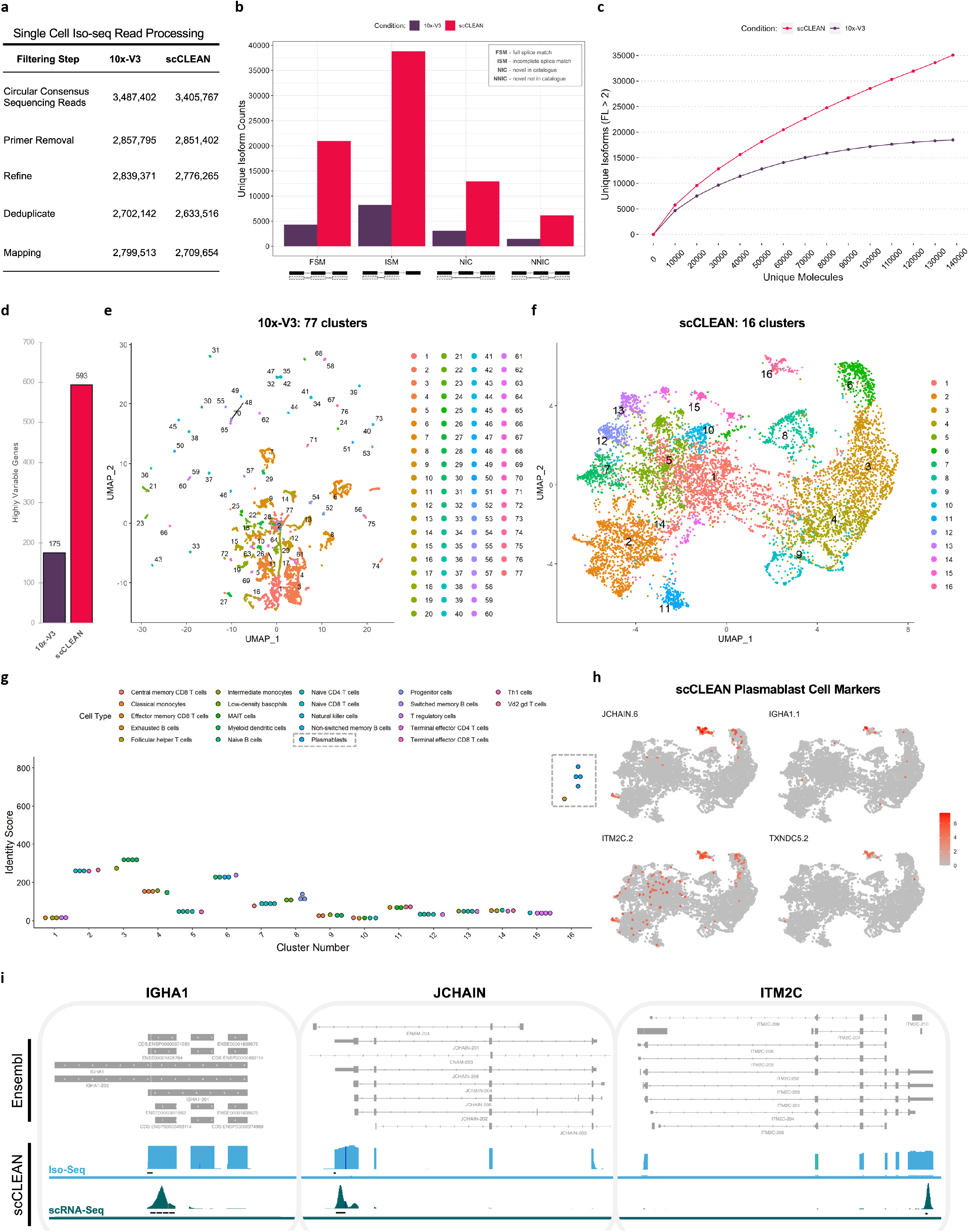
Application of scCLEAN on PBMC single cell iso-seq enables clear biological interpretation. **a**, Long read sequencing read filtration workflow. **b**, Isoform classification metrics comparing 10x-V3 (purple) and scCLEAN (red). Out of the 17,131 unique isoforms identified in the 10x-V3 condition, 4,575 were novel. Whereas within scCLEAN, 78,935 unique isoforms (4.6-fold increase) were detected with 19,112 being novel (4.2-fold increase). **c**, Saturation curve of unique isoform counts. **d**, Total number of highly variable genes identified (default mean variance = 1.3) between 10x-V3 (175) and scCLEAN (593). **e-f**, Single cell clustering (UMAP) identifying 77 clusters with **e** 10x-V3 and 16 clusters with **f** scCLEAN. **g**, Cell type identification (CIPR) of the 16 scCLEAN clusters. Identity score quantifies the confidence of assigning each cluster to cell-type. Plasmablast assigned to cluster 16 boxed in grey. **h**, scCLEAN feature plots (UMAP) of isoforms from the top 4 Plasmablast cell markers according to a FACS sorted database. **i**, IGV alignment diagrams of 3 Plasmablast cell markers (IGHA1, JCHAIN, ITM2C) comparing iso-seq data (blue) and short read scRNAseq data (green) from scCLEAN depleted PBMCs.

The increase in isoforms used for downstream analysis resulted in an increase in both the selection of highly variable genes (3.4-fold) (Fig. 4d and Supplementary Fig. 5d) as well as the captured variance along each PCA component (Supplementary Fig. 5e). Furthermore, scCLEAN-treated libraries drastically improved informative clustering performance (Fig. 4e-f). With scCLEAN, 16 clusters were identified, which is consistent with our short read analysis (Fig. 2d-e). In contrast, the 10x-V3 condition at the same resolution was sparsely clustered, identifying 77 unique clusters (Fig. 4e), rendering cell type identification un-interpretable.

Due to the sparsity inherent within current iso-seq methods, clustering and cell type annotation remain challenging, as evidenced by the standard 10x-V3 condition. Nevertheless, we wanted to elucidate if marker, gene-based identification of cell types within the scCLEAN condition conveyed additional insight over scRNAseq. We found that plasmablasts were annotated with the highest confidence from all cell clusters (Fig. 4g), despite being a notoriously difficult rare cell type to characterize^40^. Analysis of our results found several genes of interest (JCHAIN, ITM2C, TXNDC5, and IGHA1) that are key markers for plasmablasts and are differentially expressed in a single cluster (16) (Fig. 4h and Supplementary Fig. 5f). Interestingly, we additionally found that key plasmablast marker genes suffer from intense 3’ bias in short-read libraries (Fig. 4i), resulting in inefficient read mapping to these transcriptomic regions, further contributing to poor plasmablast characterization. Whereas the full isoform alignment facilitated the annotation of a rare plasmablast cell population solely based on marker gene selection. Without enhanced scCLEAN resolution, plasmablasts would have been unable to be distinguished.

### Application To Homogeneous Cell Populations

While we have seen scCLEAN’s applicability to improve cell clustering and identity assignment both in heterogenous cell populations of PBMCs as well as full length isoforms, we wanted to evaluate the ability of scCLEAN to identify cell lineages from populations of relatively homogenous makeup but integral in the pathogenesis of disease state. To characterize the baseline of coronary artery predisposition toward atherosclerosis^42^, ~50,000 primary vascular smooth muscle cells (VSMCs) were isolated taken from two distinct vascular sites (coronary and pulmonary arteries) representing 49% and 51% of the total cell population (Fig. 5a). Validating the applicability to diverse tissue types, initial scCLEAN performance metrics compared similarly to those observed in PBMCs, successfully redistributing 35% of genomic aligned reads, increasing the number of informative transcriptomic reads by 58%, and increasing the proportion of informative UMIs from 66% to 95% (average across 8 samples) (Supplementary Fig. 6a-g).

**Fig. 5:**
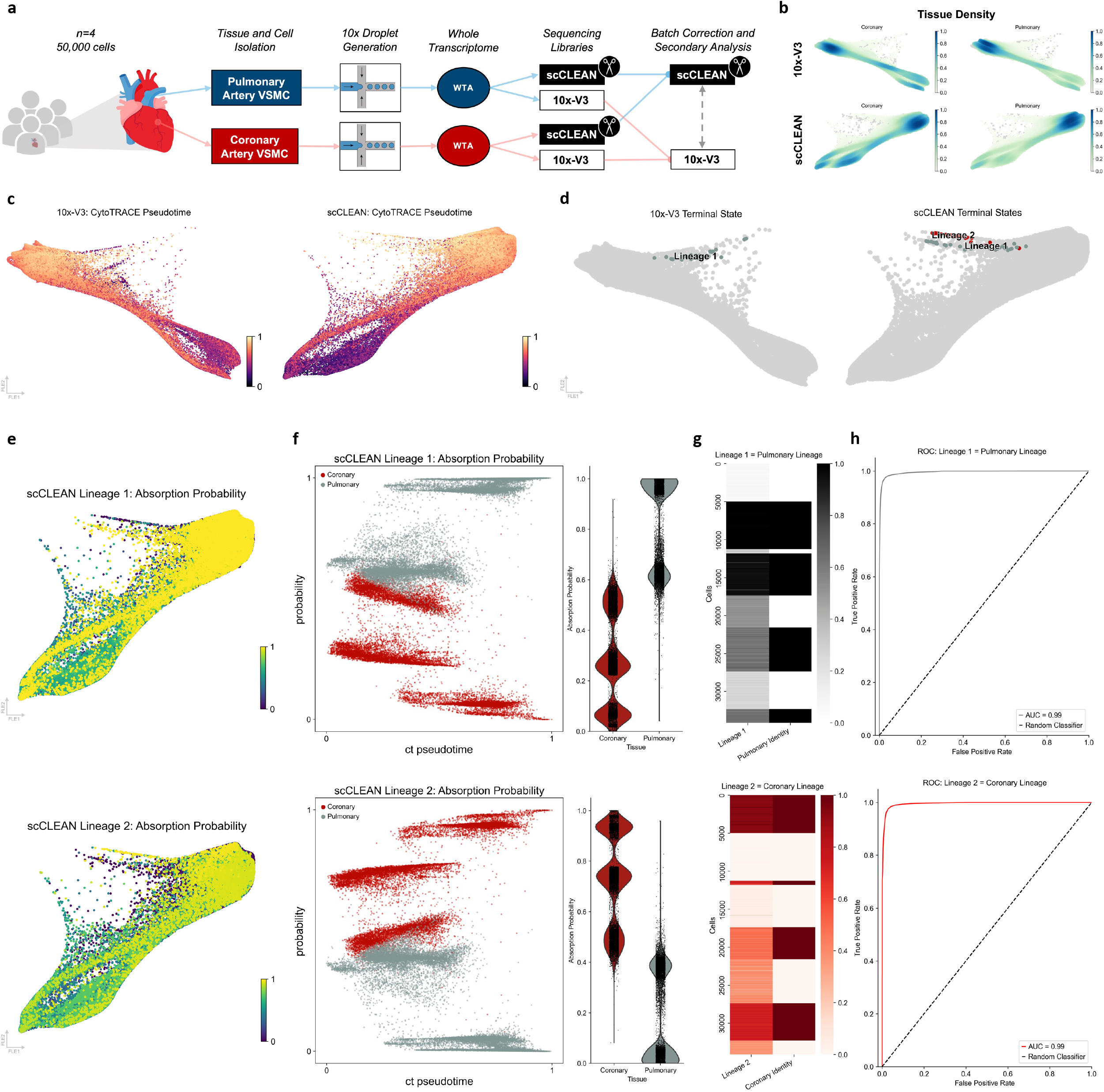
Trajectory classification of vascular smooth muscle cells with stark delineation between coronary and pulmonary artery. **a**, Schematic figure of experimental workflow. ~50,000 VSMCs were isolated from 4 patients and 2 tissue locations, coronary and pulmonary artery. Both scCLEAN and 10x-V3 represent the same population of cells. **b**, Density of cells along the force directed layout (FLE) trajectory according to their tissue of origin comparing 10x-V3 (top) with scCLEAN (bottom). **c**, Each cells transcriptional change along pseudotime (CytoTRACE) mapped onto the FLE embedding. All detected genes are utilized to automatically calculate early pseudotime (black) to late pseudotime (yellow). **d**, The total number of terminal states identified using Cell Rank comparing 10x-V3 (left) (1 lineage) with scCLEAN (right) (2 lineages). Optimal terminal states identified using Schur decomposition (gap in the real portion of eigenvalues) and then refined according to stationary distance of the coarse-grained Markov transition matrix (non-zero distance). **e**, Probability of each cell belonging to each scCLEAN lineage (absorption probability) and thus differentiating along pseudotime into either of the 2 terminal states, corresponding with lineage 1 (top) and lineage 2 (bottom). Yellow represents a 100% probability of that cell belonging to that lineage. **f**, Left, scCLEAN lineage absorption probability (lineage 1 = top, lineage 2 = bottom) plotted as a function of each cells position along the differentiation trajectory (ct pseudotime). Coloring reflects whether the cell was derived from a coronary or pulmonary artery. Right, absorption probabilities compressed along pseudotime. **g**, Heat matrix illustrating the probability of each cell belonging to each lineage paired side by side with the identity of that cell belonging to each tissue. **h**, Receiver operating characteristic (ROC) depicting the classification performance of identifying each tissue to each lineage (both pulmonary and coronary lineage AUC=0.99).

Given that VSMCs transition phenotypically between many functional states^41–45^, we performed pseudotime trajectory analysis, and selected for populations interacting in transcriptional transition. Between coronary and pulmonary, the density of cells remained comparably distributed across both conditions throughout the trajectory (Fig. 5b). We next sought to discern whether the cells followed the same trajectory or bifurcated into separate lineages. One lineage was identified in the 10x-V3 condition while two lineages were identified with scCLEAN due to the distinction between two separate terminal states (Fig. 5c-d). According to the individual cell-fate potentials, 64% and 36% of cells in the scCLEAN condition associated with lineage 1 and 2, respectively (absorption probability > 0.5) (Fig. 5e).

Since there was a roughly even split between each lineage in the scCLEAN analysis, we next sought to resolve if the tissue origin, either coronary or pulmonary artery, had any concordance to each lineage state transition. After comparing tissue identity with lineage absorption (Fig. 5f-g), it was evident that indeed each lineage was tissue specific. Receiver operating characteristic analysis confirmed the classification of lineage to artery identity with an AUC of 0.99 for both lineages (Fig. 5h). To validate correct lineage analysis and ensure the inability of the standard condition to classify tissue specificity, two lineages were forced in the 10x-V3 condition, but discriminatory power was non-existent (AUC=0.51) (Supplementary Fig. 7a-g). Consequently, by increasing signal to noise and capturing higher biological variance, scCLEAN enabled blind algorithmic classification of cell identity to lineage with outstanding accuracy, in a system that otherwise couldn’t be distinguished. Furthermore, as both populations of cells are VSMCs with only tissue origin differences, scCLEAN primes additional insights into the pathobiology of disease states.

Considering we could delineate between two tissue specific lineages and there was a strong association of genes to each lineage across pseudotime (Fig. 6a), we sought to gain a deeper understanding of how these lineages diverged by generating a principal tree with discrete branches and bifurcations (Fig. 6b). This enabled the distinction between lineage markers, genes whose pattern of expression across pseudotime are unique to either the coronary or pulmonary terminal state (Fig. 6c-e), and transition markers, genes responsible for initiating the separation (Fig. 6f). Two of the top pulmonary lineage markers were SERPINF1 (PEDF) and SFRP1 (FRZA). The complete absence of expression of these markers within the coronary lineage is paired with the up-regulation of SERPINE1 (PAI-1) and SERPINE2 (Fig. 6a).

**Fig. 6:**
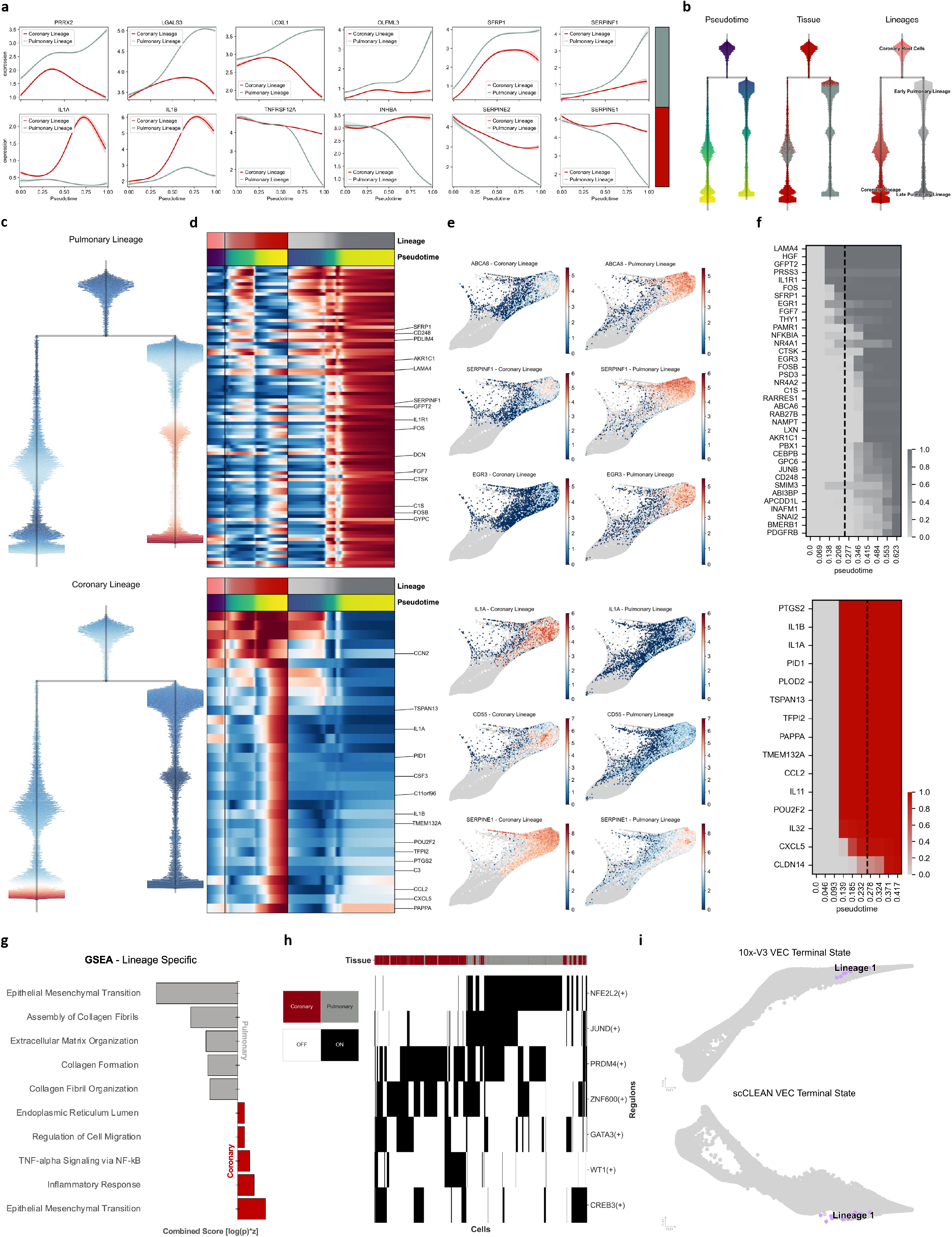
Lineage characterization reveals basal transcriptional states informing the basis of coronary cardiovascular disease. **a**, Genes with the highest correlation associated with each terminal state. Average gene expression sampled from 200 cells with a probability > 0.5 associated with each lineage (pulmonary lineage = top, coronary lineage = bottom) and mapped along pseudotime. **b**, Dendrograms representing the principal graph (scFATES) inferred from scCLEAN terminal states. Left, blue represents early pseudotime and yellow represents late pseudotime. Middle, cells colored according to tissue of origin (coronary = red, pulmonary = grey). Right, 4 distinct nodes of the principal graph labeled according to tissue proportion. **c**, Tree diagram highlighting onset of characteristic gene signatures (red) associated with pulmonary lineage (top) and coronary lineage (bottom). **d**, Gene expression matrix of highest specificity (effect>0.3) depicting early coronary root cells (left), coronary lineage (middle) (32 genes), and late pulmonary lineage (right) (91 genes). The top 15 genes according to correlation were annotated. **e**, Feature plots (FLE) comparing expression of 3 lineage markers to the pulmonary lineage (top) and 3 lineage markers to the coronary lineage (bottom). **f**, Genes responsible for pulmonary lineage (top) and coronary lineage (bottom) bifurcation in order to reflect early decision-making processes prior to separation. Matrix contains inclusion timing for each gene in each probabilistic lineage projection. **g**, Gene set enrichment analysis (GSEA) determined from all lineage specific genes with a correlation greater than 0.3. Combined score incorporates gene overlap and statistical significance. Coronary lineage genes uniquely overlap with inflammatory response and TNF-alpha signaling. **h**, Regulon activation per cell (SCENIC) (black = on, white = off) calculated from the top 5,000 coronary lineage cells and the top 5,000 pulmonary lineage cells. The top transcription factors ranked according to the average difference between lineages. NFE2L2 turned on in 91% of cells derived from the pulmonary lineage but only 1% of cells derived from the coronary lineage. **i**, Vascular endothelial cell (VEC) trajectories (FLE) representing ~30,000 cells and 2 donors. Comparison of terminal states between 10x-V3 (top) and scCLEAN (bottom).

Providing evidence for accurate biological distinction, the top coronary transition markers (IL1A, IL1B, IL11, IL32, CCL2, CXCL5) are inflammatory markers (Fig. 6f). Gene-set enrichment analysis (GSEA) further confirmed the coronary specific inflammatory signatures, with considerable overlap with inflammatory response and TNF-alpha signaling via NF-kB (Fig. 6g). Based on the observation of distinct transcript expression patterns seen so far, we sought to identify regulatory mechanisms associated with each coronary and pulmonary lineage to inform future exploration and potential avenues for therapeutic intervention. We found that nuclear factor erythroid 2-related factor 2 (NFE2L2) was activated in 91% of cells in the pulmonary lineage but only 1% of cells in the coronary lineage (Fig. 6h).

While scCLEAN enables novel lineage distinction in VSMCs with vital biological relevance, we sought to understand if similar results could be produced on a cohort of ~30,000 primary vascular endothelial cells (VECs) derived from coronary and pulmonary arteries from two donors. As VSMCs and VECs are known to communicate to promote proper vessel wall formation and function^48^, it is plausible to suggest that coronary and pulmonary VECs would also have stark differences in gene expression profiling contributing to risk of the diseased state. Proceeding with trajectory inference in VECs, we surprisingly only classified a single terminal state of VECs under both 10x-V3 and scCLEAN conditions (Fig. 6i) suggesting that VECs do not diverge to distinguishable lineages. This gives further credence that the divergent qualities found in VSMCs with scCLEAN are not due to technical or computational artifact.

## Discussion

Herein we detail scCLEAN, a novel methodology to improve the performance of scRNAseq by removing highly abundant, uninformative molecules and subsequently transferring reads to lowly expressed genes *in vitro*. Instead of 0.7% of genes comprising 45% of transcriptomic reads in standard samples, scCLEAN harnesses CRISPR/Cas9 enabling 99.3% of genes to encompass 94% of transcriptomic reads. This complete overhaul in read composition translates to a significant boost in signal to noise, increasing the biological variance incorporated into the latent representation as validated in four distinct cellular applications. Overall, the benefits surpass the negligible offtarget effects representing 0.06% of the transcriptome.

Previous approaches to combat molecular dropout typically revolve around identifying relevant genes of interest and targeting them with some form of capture-based approach. While different iterations have been successfully applied to increase transcription factor resolution^46^, antigen receptor sensitivity^47^, and combinatorial perturbation scale^48,49^, the gene panel size is currently limited to roughly 1,500 genes, less than 5% of the transcriptome (1,500/36,601) (10x Genomic Reference). Consequently, only genes known a priori to be biologically relevant are likely to be considered, thus eliminating, novel unbiased identification of gene associations. Conversely, scCLEAN leverages 99% of the transcriptome. Furthermore, while CRISPR/Cas9 has been previously integrated into multiple molecular methods to remove unwanted sequences^50–53^, scCLEAN drastically expands the scope of the application, enabling the enhancement of biological signal and downstream analysis. In addition, scCLEAN is the first application designed to remove multiple uninformative targets (not limited to rRNA) from polyA-captured scRNAseq libraries.

The power of scRNAseq analysis lies in the ability to identify both large and minute contributions to the biology of the end result observed in health or disease. By improving carryover of the cellular heterogeneity incorporated into downstream analysis, the confidence and statistical power of biological inference is enhanced. The improved resolution enables nuanced transcriptomic distinction such as the characterization of rare cell types in PBMCs, the identification of cellular states through enhanced isoform detection, and the association of novel genetic signatures to coronary and pulmonary lineages within VSMCs. While these tissue specific lineages share considerable phenotypic similarity and are evenly distributed throughout the trajectory, it is comprehensible that there be some fundamental distinction considering pulmonary VSMCs are derived from the neural crest and coronary VSMCs are derived from the epicardium^54^. To the best of our knowledge, the results presented above characterize for the first time pre-symptomatic, basal level differences in how the two tissue lineages diverge transcriptionally, setting up a stark juxtaposition of relatively similar vascular beds.

The ability to distinguish between coronary and pulmonary VSMC plasticity could be instrumental in the understanding of underlying predisposition to disease states (e.g. atherosclerosis), onset, and/or progression. Reassuringly, scCLEAN provides insights that are aligned with the field’s current disease characterization and novel discoveries. Coronary artery specific inflammation, as validated by the top coronary lineage markers as well as GSEA, is considered a hallmark process of atherogenesis^55^. Furthermore, SERPINF1 (PEDF) and SFRP1 (FRZA), top pulmonary lineage markers, exert anti-proliferative effects on smooth muscle cells both *in vitro* and *in vivo*, arresting cell-cycle progression^56,57^. These findings coincide with the up-regulation of SERPINE1 (PAI-1) and SERPINE2, top coronary lineage markers, which promote VSMC proliferation, migration, and vascular remodeling^58,59^. These findings allude to higher phenotypic switching from contractile to a proliferative state within coronary cells, laying the groundwork for the aberrant VSMC proliferative phenotype observed within atherosclerosis^54^.

In addition to validating the biological relevance of each tissue lineage, scCLEAN facilitated original insights through regulon analysis. NFE2L2, which was exclusively identified in the pulmonary lineage, is pivotal in the regulation of antioxidant proteins that protect from injury and inflammatory induced oxidative damage^60^. This finding coincides with deficient SERPINF1 (PEDF) expression in the coronary lineage, since SERPINF1 quenches oxidative stress^61^. Considering oxidative stress contributes to the development of vascular diseases such as atherosclerosis^60^, the lack of NFE2L2 regulon activation in coronary cells may provide insight into early disease onset and warrants further investigation. Consequently, we further bring to light a novel distinction between the ability of each tissue to respond to oxidative stress, motivating the potential development of cell-mediating therapeutics to prevent the onset of atherosclerosis. Due to the ability of scCLEAN to derive biological insights previously obscured in noise, we envision widespread incorporation into existing high-throughput single-cell techniques.

Potential limitations unaddressed in the scope of this study include the relative importance of the 255 genes removed with scCLEAN. For example, if aberrances in ribosomal gene expression are important for cancer specification in the sample type of interest due to a dysregulation in cell cycling^62^, the 90 ribosomal genes targeted in scCLEAN should not be included. In addition, while the 155 non-variable genes also targeted for removal were determined to be uninformative for cell state determination within 14 sample types, it is noted that they still in-part serve some context-dependent biological role. Furthermore, although one of the main benefits of scCLEAN is the flexibility to apply this method to any scRNAseq platform, the relative abundance of all targeted regions might vary according to method, potentially altering scCLEAN depletion performance and sensitivity improvement. For example, if a method has poor capture sensitivity and generates low complexity libraries, then scCLEAN has a higher probability of replacing highly abundant molecules with PCR duplicates, providing limited benefit. However, a large benefit in the methodology of scCLEAN is that it is customizable to fit a desired platform of interest, as well as target a desired gene cohort of disinterest.

As evidenced by the easy transference of scCLEAN to enhance single-cell isoform detection, scCLEAN holds substantial potential to improve single-cell methodologies beyond RNA-seq, such as spatial transcriptomic assays that utilize NGS spatial barcoding^63–67^. In addition, we expect scCLEAN to impact single-cell multi-omic sequencing technologies, considering all existing combinations contain a transcriptomic component^68^. While providing comprehensive insights, combining modalities such as transcriptome, genome, epigenome, and proteome from a single cell only exacerbates the noise contributed by each technique. scCLEAN provides a unique opportunity to address these additional bottlenecks and universally improve resolution.

## Methods

### sgRNA Design and Generation

We curated a set of 14 different scRNAseq datasets from SRA for various human sample types. Note that samples in each of these datasets were prepared in accordance with the 3’ 10x Chromium V3 technology (10X Genomics, Pleasanton, CA).

### Genomic Intervals

While an average of 93% of the reads from these datasets aligned to the genome, only 61% (average) of total reads aligned to the transcriptome, and as single-cell feature matrix files only compose of transcriptomic alignments, theoretically, 32% of reads are completely ignored in analyses and are ideal candidates for removal. For each sample, reads were aligned to a reference index built from Ensembl transcriptome (Homo_sapiens.GRCh38.cdna.all.fa) using the flag --twopassMode Basic to maximize the read alignment, revealing conserved intervals that can be targeted for removal. All reads that overlapped with a transcript interval were removed. The filtered bam files from each tissue were then used to calculate the coverage over 500bp intervals across the human genome using bedtools coverage, with assessment that no transcriptomic regions are targeted.

scCLEAN guides that target genomic but not transcriptomic intervals consist of a combination of two cohorts: 500bp intervals with the highest shared coverage across all 14 tissues, and the combination of the top 500bp tissue specific intervals. Interactive Genomic Viewer (IGV)^69^ was used to observe the interval location across tissues relative to the transcriptome and genome. Intervals fall into 3 distinct categories: intergenic regions, unannotated genomic deserts, and regions that are alongside annotated transcriptomic intervals.

### Non-Variable Genes (NVG)

To identify a set of highly expressed protein coding genes that did not play a significant role in identifying cell state or type across a diverse set of human tissues, all 14 SRA datasets were aligned to the 10x transcriptome (refdata-gex-GRCh38-2020-A) using cellranger count and underwent standard secondary processing according to Pegasus (https://github.com/lilab-bcb/pegasus)^70^. For each dataset, the top 5,000 highly variable genes were selected. A list of all remaining genes that were not the top 5,000 highly variable were then recorded for each dataset. Conserved genes that were not the top 5,000 and shared across all 14 datasets were then reviewed and the 155 NVG were then selected according to the rank of average expression across all datasets. The NVG were assessed for identity consistent with primarily housekeeping genes with PanglaoDB^71^. 73% of the 155 NVG directly overlapped with the top 1,000 genes according to the Ubiquitousness Index. In fact, for all 255 protein coding genes targeted with scCLEAN, 79% are within the top 1,000 most ubiquitous genes.

## PBMC Samples

### Sample Extraction

Blood was obtained from the Scripps Normal Blood Donor Service from anonymized donors and fully compliant with approved IRB protocols for human subjects. Blood was collected and prepared for downstream single-cell applications (see Supplementary Methods).

### Single-cell Library Generation

The cell suspension was then prepared for droplet generation according to the manufacturer’s suggested protocol (10X Genomics, CG00053 Rev C). Four channels aiming to recover 10,000 cells were loaded on the 10x Chromium Controller and the protocol was followed according to the manual Chromium Next GEM Single Cell 3’ Reagent Kits v3.1 (Dual Index) (CG000315 Rev C). Using the Chromium Next GEM Single Cell 3’ Kit v3.1 (16 rxns PN-1000268), all four libraries were generated and carried forward. 120ng of cDNA from each channel was carried forward into step 3, through step 3.4, post ligation cleanup. Three libraries underwent scCLEAN depletion (see scCLEAN Depletion) while the control followed the standard 10x protocol. Both conditions underwent the same number of PCR cycles for single index sample PCR and were sequenced on a Nextseq 2000 (Illumina, La Jolla, CA) with the following sequencing schematic: R1 - 28, I1 - 8, R2 - 91. For all downstream analysis, all four libraries were down sampled to the exact same read depth of 310 million reads (~27k reads/cell) using seqtk with the appropriate flags.

### scCLEAN Depletion

The Chromium Next GEM Single Cell 3’ protocol v3.1 was followed per manufacturer’s suggested protocol until the end of Step 3.4 – Post Ligation Cleanup – SPRIselect. Immediately after Step 3.4, the scCLEAN depletion protocol was followed (detailed protocol see Supplemental Methods). Briefly, the library is eluted in 16 μL of CRISPRclean Nuclease Free Water. Next, 15ul of the library is then carried forward into Cas9-RNP incubation with the components in CRISPRclean Single Cell RNA Boost Kit (Jumpcode Genomics KIT1018, San Diego, CA). For each reaction, a RNP consisting of 1.0ul 10x Cas9 buffer, 1.0ul RNAse inhibitor, 3.9ul of 1ug/ul sgRNA library, and 2.3ul of Cas9 enzyme is mixed and left at room temperature for 10 minutes. The RNP, library from step 3.4, as well as an additional 1.5ul of 10x Cas9 buffer is mixed and incubated at 37°C for 60 minutes (total reaction volume ~25ul). After incubation, a 0.6x SPRI clean is performed using AMPure XP beads (Beckman Coulter A63881) to remove the cleaved fragments. The cleaned library is then eluted in 30ul of Nuclease-Free Water and the Chromium Next GEM Single Cell 3’ protocol v3.1 at Step 3.5 – Sample Index PCR was followed until completion of the libraries.

## scCLEAN Performance Evaluation

### QC Comparison

To properly compare the change in library composition due to scCLEAN, both the 10x-V3 control condition as well as the scCLEAN depleted condition were first down sampled to the exact same read depth using ‘seqtk sample’ with the appropriate flags. Next, they were each aligned to two separate indexes using ‘cellranger count.’ One index was the standard 10x transcriptome (refdata_gex_GRCh38_2020_A), and the second index was the same exact transcriptome except the index was built using a gtf file where the annotations associated with the 255 targeted protein coding genes were removed (Supplementary Methods). The filtered matrix files used for downstream processing were exclusively taken from the alignment to the full, not-masked, standard transcriptome. All statistical comparisons between the distribution of quality metrics were performed using the Mann-Whitney U rank test with the Benjamini-Hochberg correction with default thresholds. See supplementary methods for read-redistribution calculation.

## PBMC Secondary Analysis

### Unsupervised clustering

Cell clustering was performed with Seurat v.4.1.1 R toolkit^3^. Genes expressed in less than 3 cells were filtered from the data and cells with less than 200 genes and 500 UMIs. Additionally, cells in the 99th percentile of UMIs were filtered to remove homotypic doublet influence. The distribution of MT genes as a percentage of total contributing features was examined and cells within the top 5th percentile were removed. The “SCTransform” function was performed to normalize the count data prior to principal component analysis^72^. We used a residual variance cutoff of 1.3 to calculate the optimal number of variable features. Additionally, the percentage of MT genes and cell cycle scoring was regressed out in a second non-regularized linear regression using the argument “vars.to.regress.” The “RunPCA” function was used for principal component analysis using default settings and the first 50 PCs. The “FindNeighbors” and “RunUMAP” functions were used using default settings and dims parameter set to 30. The first 30 PCs were used to construct the shared-nearest neighbor (SNN) network graph followed by construction of the Uniform Manifold Approximation and Projection (UMAP) dimensionality reduction^73^. Clusters were identified using the “FindClusters” function using the Louvain algorithm with multilevel refinement and resolution set to 1.2. This resolution was used to first over-cluster the data for subsequent doublet removal.

### Doublet Removal

The R package “scDblFinder” was used for the identification of heterotypic doublets using a cluster-based generation of artificial doublets^74^. Because “SCTransform” uses a binomial regression model for UMAP plot generation and 2D visualization, we used the “NormalizeData” function in Seurat to log-normalize the counts data for artificial doublet creation before doublet removal. Using the “scDblFinder” function, we set the cluster assignment to those calculated from Seurat v4.1. Additionally, we set the “nFeatures” parameter equal to the number of variable features calculated from the residual variance cutoff of “SCTransform.” We also set the argument “processing = normFeatures” to use normalized counts of genes in calculation of doublets. Additionally, the arguments “dims = 30” and “PCs=30” were used for the number of dimensions and principal components to be used, respectively. Once doublets were removed, we repeated the unsupervised clustering as described above.

### Clustering Resolution Selection

Semi-unsupervised/unbiased techniques for cell clustering were also utilized. Optimal clustering resolution for 2D visualization comparison between the control and scCLEAN conditions was determined using ROGUE R package^75^. The average ROGUE statistic across all cell clusters at a range of clustering resolutions (0.1 – 1.2) was first calculated in the standard condition. The optimal clustering resolution of 0.4 was chosen because the ROGUE statistic reached a maximum saturation of cell cluster purity at this given resolution (Supplementary Fig. 1b). For consistency and direct comparison of cell clustering efficiency, we also set the clustering resolution at 0.4 in the scCLEAN condition.

### Cell-cell communication inference

The CellChat R toolkit was used to infer inter- and intra-cellular communications^76^. Each sample in its entirety was processed with default thresholds according to the tutorial. Next, the control and depleted conditions were subsetted to include cell clusters that are consistent in both cell numbers and cell-type annotations across both conditions. As a result, a total of 9 cell types were chosen to compare between conditions: CD4+ Memory T, Naïve Memory T, Classical Mono, Mature NK, MAIT, MEP, Naïve B, Memory B, and pDC. Additionally, geometric sketching using the geosketch package was employed to down sample each condition to 8,000 cells^77^. We followed the workflow as outlined in the CellChat tutorial using default settings with Communication Probability population size = TRUE.

### Cluster marker analysis and cell type annotation

Cell types were annotated with a combination of the R package CIPR (v.0.1.0), which leverages reference cell populations and complex multi-gene expression signatures, and canonical immune cell markers^78^. We first performed DEG analysis using the “FindAllMarkers” function in Seurat R package. The log-normalized gene expression counts were used as input and genes were quantified as upregulated with a log2 foldchange > 1 and adjusted p-value < 0.01. The output of “FindAllMarkers” was used as input into the CIPR package for cluster scoring and annotation. The reference database used was “hsrnaseq” which is a sorted human RNAseq dataset^79^. Additionally, some immune cell subtypes could not be effectively identified from CIPR. Thus, we employed the use of classical markers to annotate cell sub-types and verify CIPR annotated cell clusters. See supplementary methods for a full list of cell type identification markers.

### Differential Expressed Genes (DEGs)

We performed DEG analysis using the “FindMarkers” function in Seurat R package. We compared the log-normalized expression values between 10x-V3 cell populations and scCLEAN depleted cell populations, respectively. The fold changes of the mean log-normalized expression values between the two cell populations were calculated. Only genes which were expressed in at least 10% of cells for each condition were considered. Genes significantly enriched had a Log2 fold change threshold of 0.25 and have an adjusted p-value < 0.05.

## Orthogonal Validation

### Random Matrix Theory – Signal to Noise

For each sample, random matrix theory was used to distinguish between signal and noise^80^, and consequently compare between scCLEAN and 10x-V3. The python tool randomly (https://github.com/RabadanLab/randomly) was used with minimal filtering to accurately characterize the effect of scCLEAN on the signal to noise ratio. Data processing followed the randomly tutorial, initializing the randomly.Rm() model, and using the function preprocess with the following arguments: min_tp=1, min_genes_per_cell=1, min_cells_per_gene=1, refined=True. The model was then refined using the command refining with the flag min_trans_per_gene=1.

### Deep Learning Unsupervised Clustering

To ensure that the identification of additional cell clusters with scCLEAN was not an artifact of hyperparameter tuning, deep embedding for single-cell clustering (DESC) (https://github.com/eleozzr/desc) was used, leveraging deep learning to identify cell clusters in an unsupervised fashion, and assign a confidence score of assigning each cell to a specific cluster following similar initial filtering described above^81^. Despite being an unsupervised method, the proper clustering resolution still had to be manually chosen, so to compare DESC outputs between scCLEAN and 10x-V3, a range of Louvain resolutions were selected (0.1, 0.2, 0.4, 0.6, 0.8, 1.0, 1.2, 1.4, 1.6, 1.8, 2.0). For each resolution x, the model was trained using the function train with the following arguments: dims=[anndata.shape[1], 128, 32] tol=0.001, n_neighbors=10, batch_size=256, louvain_resolution=x, do_tsne=True, learning_rate=300, do_umap=True, num_Cores_tsne=4.

### Gene Mapping onto the Latent Space

SIMBA (1.1) (https://github.com/pinellolab/simba) was used to embed individual genes alongside cells in feature space, enabling the characterization of cell type-gene specificity^82^. Cellranger output files filtered_feature_bc_matrix.h5 for each condition were input and consequently subset to only the cells that remained post filtering and were used for downstream analysis (see PBMC Secondary Analysis). The matrices were then processed according to the SIMBA tutorial and documentation. Data was then discretized using tl.discretize with n_bins set to 5. The graph was then generated with use_highly_variable set to True. The parameters to train the model were left on default settings. Cell type annotations were lifted over from the Seurat analysis (see Cluster marker analysis and cell type annotation). SIMBA UMAP plots were generated with n_neighbors=20, n_components=2, and random_state=10. All statistical comparisons between the distribution of specificity metrics were performed using the Mann-Whitney U rank test with the Benjamini-Hochberg correction with default thresholds.

To illustrate the additional gene-cell type specificity in the scCLEAN versus 10x-V3 condition, SIMBA was used to quantify genes within a certain radius of the densest point of a cell cluster. Within the scCLEAN condition, each of the four additional cell types that were identified were isolated and the mean x and y coordinates in UMAP space were calculated. If the cell type consisted of more than one cluster, the cluster with the largest density of cells was selected. From the central point, genes were queried using tl.query with use_radius=True and r=0.5 to identify genes within a radius of 0.5 in UMAP coordinate space. Considering, in the 10x-V3 condition the four cell types were not identified, it is infeasible to directly calculate the center of the highest density cells pertaining to that cell type. As a proxy, 3 out of the 4 clusters could be visually distinguished in UMAP space due to the surrounding cell types as well as physical characteristics such as distensions of cells. For example, in the scCLEAN condition, an entirely isolated island of cells consists of two cell types, Mature Natural Killer cells and S100B+ Natural Killer cells. In the 10x-V3 condition, the same island exists, but only consists of Mature Natural Killer cells. Consequently, the radius in the 10x-V3 condition was selected to closely approximate the location of the S100B+ Natural Killer cells in the corresponding scCLEAN condition. Proliferating Natural Killer cells were the only cell cluster that could not be identified since an additional isolated cluster did not exist in the 10x-V3 condition. Consequently, comparative analysis performed for the four cell types uniquely found within the scCLEAN condition were qualitative and should be interpreted for intuitive purposes exclusively.

## Single-Cell Iso-Seq

PBMC samples were isolated from a donor and prepared using 10X Genomics Chromium Next GEM Single Cell 3’ Reagent Kit (v3.1) protocol (10X Genomics, CG000315 Rev C) as described above. 100ng input of GEM barcoded WTA (cDNA) was then treated with CRISPRclean Cas9/Single Cell Boost Guide RNA complexes for cleavage of targeted sequences. The ribonucleoprotein complex reaction was adjusted to target <2kb single-cell cDNA short transcripts, ensuring a 1:200:500 molar ratio of target to enzyme to guide. After CRISPR digestion, the cleaved targeted sequences were 1X magnetic bead-based size selected using the ProNex Purification System. The remaining desired un-cleaved cDNA samples were PCR reamplified alongside standard untreated cDNA samples from the same GEM channel cell population. The amplified cDNA samples were then 0.95X ProNex size selected for desired <2 kb short transcripts.

Following single-cell cDNA quantitative mass and qualitative size validation for desired small transcripts, SMRTbell libraries were generated following the “Single-Cell Iso-Seq SMRTbell Express Template Prep Kit 2.0 Procedure” as per the manufacturer’s protocol (PacBio PN 101-892-000 v1, Menlo Park, CA). Each library was sequenced individually on the PacBio Sequel II System targeting ~3 million full-length single-cell transcript reads per SMRT Cell 8M.

### Generation of CCS reads, Barcode Correction, and Deduplication

Circular Consensus Sequencing (CCS) reads were generated from subreads files using the CCS bioconda package v6.4.0 with default parameters. Both CCS files were down sampled to 3.4 million reads using samtools v1.12 view command using the –s parameter set appropriately. Primers were removed using Lima v2.5.0 (–isoseq, –peek-guess) using the 10X Chromium Next GEM Single Cell 3’ (10X GEX 3’) primers as inputs. To identify and clip both UMIs and cell barcodes, the isoseq3 v3.7.0 tag command was used with the design parameter set to “T-12U-16B” to match the 10X GEX 3’ barcode schematic. Poly A selection and concatemer removal were performed using isoseq3 refine.

Barcode correction was performed using the isoseq3 correct command using the 10X barcode whitelist (3M-february-2018.txt) obtained from the Cell Ranger v6.2.0 package. The refined, corrected reads were then sorted by corrected barcode using samtools (sort -t CB) to prepare for deduplication. We performed deduplication using the isoseq3 groupdedup command.

### Alignment of Deduplicated Reads to Genome

Deduplicated reads were aligned to the RefSeq genome (GCF_000001405.39_GRCh38.p13_genomic.fa)^83^ using pbmm2 v1.8.0. pbmm2 was invoked using the parameters –preset ISOSEQ and –sort. To generate a gtf file containing unique isoforms, we used isoseq3 collapse on the mapped, deduplicated reads.

### Isoform Classification

Isoform classification was performed using the PacBio bioconda pigeon package v0.1.0. The RefSeq gtf and fasta file were indexed for compatibility with pigeon, using the pigeon index and samtools faidx commands respectively. Isoform classifications were obtained using the pigeon classify command with the –fl parameter to append unique molecule count to facilitate downstream filtering. Classified isoforms were then filtered for intra-priming and RT-switching using the pigeon filter command. We applied an additional filter by removing any isoform with less than 2 associated unique molecules (FL read column > 1).

Exclusively to calculate the boost in informative isoforms, isoforms corresponding to the 255 genes that were targeted by scCLEAN were removed from the total isoform metrics (Fig. 4b-c). For all downstream biological processing, all isoforms were retained. Seurat compatible inputs were generated using the final, filtered isoforms (containing all genes) using the pigeon make-seurat command.

### Single-Cell Iso-Seq Analysis

Cell clustering was performed with Seurat v.4.1.1 R toolkit with genes expressed in less than 3 cells and cells with less than 10 genes filtered from the data. Additionally, as an initial pass of low-quality cells filtering, those with less than 25 UMI’s detected were filtered from the data. The “SCTransform” function was performed to normalize read depth prior to principal component analysis. We used a residual variance cutoff of 1.3 to calculate the optimal number of variable features. The “RunPCA” function was used for principal component analysis using default settings and the first 50 PCs. The “FindNeighbors” and “RunUMAP” functions were ran using default settings and dims parameter set to 30. The first 30 PCs were used to construct the shared-nearest neighbor (SNN) network graph followed by construction of the Uniform Manifold Approximation and Projection (UMAP) dimensionality reduction. Clusters were identified using the “FindClusters” function using the Louvain algorithm with multilevel refinement and setting the maximum resolution to 0.6. This resolution was used to first overcluster the data for subsequent doublet removal (see doublet detection). In a second pass of low-quality filtering, we filtered low-quality cells with low gene counts, relative to UMI counts (i.e., red blood cells). This was done by filtering cells in the bottom 0.01 quantile for gene counts to account for cells that pass the >25 UMI threshold but did not have the same relative proportion of unique gene counts. A second round of semi-supervised clustering was performed as previously described. The “FindClusters” function using the Louvain algorithm with a sequential number of clustering resolutions from 0.05 to 0.6 at 0.05 intervals. A final clustering resolution of 0.35 was subsequently chosen and applied to both the “10X-V3” and “scCLEAN” conditions. For cell annotation, we used a sorted scRNAseq dataset to identify the most significant genes expressed in plasmablasts (Log2 fold change >4 and FDR < 0.005)^79^, and CIPR was utilized with default thresholds to calculate identity score.

## Cardiovascular Samples

### Sample extraction and single-cell isolation

Human coronary and pulmonary artery smooth muscle cells from healthy donors were obtained from Cell Applications (San Diego, CA). Specifically, cells were matched to the same original donor and utilized for scRNAseq at low passage (less than passage 3). A total of four distinct individual donors were used. Cells were cultured in growth medium (Cell Applications) at 37 °C with 5% CO2 atmosphere in T25 flasks (Corning, Corning, NY) and were detached from cell culture flasks using Trypsin (Invitrogen) incubation at 37 °C for single-cell suspension. Cell viability was assessed using trypan blue staining (Invitrogen, Waltham, MA) and Countess II (Invitrogen) with samples having viability of >90% being used for scRNAseq.

Human coronary and pulmonary artery vascular endothelial cells from healthy donors were obtained from Cell Applications (San Diego, CA) and prepared identically to VSMCs as detailed above. Except for VEC samples, two distinct individual donors were utilized, and growth medium was specific to endothelial cells (Cell Applications). Both VEC and VSMC samples then underwent scRNAseq and scCLEAN protocols (see PBMC Samples). Sequencing libraries were generated on a NovaSeq 6000 XP workflow (Illumina) with the following read parameters: R1-28, R2-92, I1-8, I2-8.

### Single-cell analysis

For every biological sample, a scCLEAN library and a 10x-V3 library were generated, representing the exact same cell population. Each tissue varied in sequencing depth, but both conditions for each tissue were down sampled to the exact same sequencing depth using ‘seqtk sample.’ For secondary analysis, a cell ranger index was generated using Ensembl sequence file Homo_sapiens.GRCh38.104.fasta and an annotation file using ‘cellranger mkref’ with the accompanying Ensembl Homo_sapiens.GRCh38.104.chr.gtf with the flag -- attribute=gene_biotype:protein_coding. The samples were then aligned using ‘cellranger count’ with default parameters.

Each sample UMI count matrix next underwent quality control filtering, removing cells with fewer then 200 genes and 500 UMIs. In addition, the cells representing the top 1% of UMI counts were removed as well as genes that were found in fewer than 3 cells. A MT threshold was set using the python implementation of miQC (see Supplementary Methods “Python miQC Filtering”) with a threshold adjustment of 10 median absolute deviations (see Supplementary Methods). All samples from each condition were then aggregated into one anndata object using Pegasus (1.5.0) command ‘pegasus aggregate_matrix.’ Initial processing was performed according to the Pegasus analysis tutorial with default parameters. Doublets were inferred and removed using Pegasus ‘infer_doublets’ function. with the following arguments: channel_attr=‘Channel’, clust_attr=‘louvain_labels’, expected_doublet_rate=0.001. ~7.5% of the total cells were predicted as doublets and were removed. After filtration, initial processing was repeated, and batch correction was performed using run_harmony with default parameters^84^. The Louvain clustering was calculated with the resolution 0.4. The diffusion map and force directed layout graph was generated using the function diffmap and fle, respectively. Identified clusters that were not connected to the main trajectory according to the force directed layout graph were removed from downstream analysis.

### Trajectory Inference

Cell Rank (https://github.com/theislab/cellrank)^85^ was used to identify trajectories. The CytoTRACEkernel was used to generate cell differentiation pseudotime^86^. The transition matrix was calculated using the function ‘compute_transition_matrix’ with the flags threshold_scheme=‘soft’ and nu=0.3. To compute terminal states, and the associated fate probabilities, the Generalized Perron Cluster Cluster Analysis (GPCCA) estimator was utilized^87^. To identify the top terminal states, schur decomposition was performed using the function compute_schur with n_components=25 and alpha=0.2. For both VSMC sample conditions, an eigengap was detected after 2 eigenvalues, so two macrostates (terminal states) were computed using compute_macrostates with n_states=2. Only the macrostates that had non-zero stationary distance (stationary distribution of coarse-grained transition matrix) were carried forward in analysis. Terminal states were established from the two macrostates, and fate probabilities were identified using compute_absorption_probabilities with time_to_absorption=‘all’. The top genes associated with each terminal state were identified using the function compute_lineage_drivers.

While Cell Rank was used to identify distinct terminal states, scFATES (0.8.0) (https://github.com/LouisFaure/scFates) was then utilized to characterize gene-lineage associations with respect to pseudotime, enabling the distinction between genes that drove the split between the two lineages as well as genes highly correlated with each^88^. The tutorial “Conversion from CellRank pipeline” was closely followed. The cell rank analysis was converted to a lineage tree using cellrank_to_tree with the flags time=‘pseudotime’, Nodes=10, and seed=10. The single tip of the tree that was identified closest to the cells of earliest pseudotime was established as the root. scFATES principal graph was calculated using tl.pseudotime with n_map=100 and seed=42. The dendrogram was established using the function tl.dendrogram with default parameters. Genes significantly associated to each section of the principal tree were calculated using test_association and fitted to the trajectory using tl.fit. Genes associated with the Coronary and Pulmonary branch were identified using tl.test_fork with the root_milestones flag set to the “Coronary Root Cells” and the milestones flag set to the two terminal branches “Coronary Lineage” and “Late Pulmonary Lineage.” Markers were selected using the function branch_specific with the same root and milestones selected as above with effect=0.3. 91 genes were associated with the Late Pulmonary Lineage and 32 genes were associated with the Coronary Lineage. To identify gene’s specific to each module (lineage) at the point of bifurcation with respect to pseudotime inclusion, the function tl.module_inclusion was used with the same root and milestones selected as above in addition to n_map=50 and parallel_mode=‘mappings.’ See supplementary methods for GSEA and regulon analysis.

## Competing Interests

At the time of manuscript preparation, J.B., D.D., A.C., K.C., J.D., A.S., S.R., K.B., and J.A. were employees of Jumpcode Genomics. All other authors declare no competing interests.

## Acknowledgements

This study was supported by the National Center for Advancing Translational Sciences, National Institutes of Health, through UL1TR002550 (EJT) as well as linked award KL2TR002552 (ACP). It was also supported by American Heart Association Career Development Award 20CDA35310187 (ACP). The content is solely the responsibility of the authors and does not necessarily represent the official views of the NIH/AHA.

## Data and Code Availability

All single-cell sequencing reads, and count matrices are in the process of being submitted to the Gene Expression Omnibus and will be fully available at a later time. All trajectory analysis code is being organized and will be available on GitHub soon.

---

**Supplementary Fig. 1:**
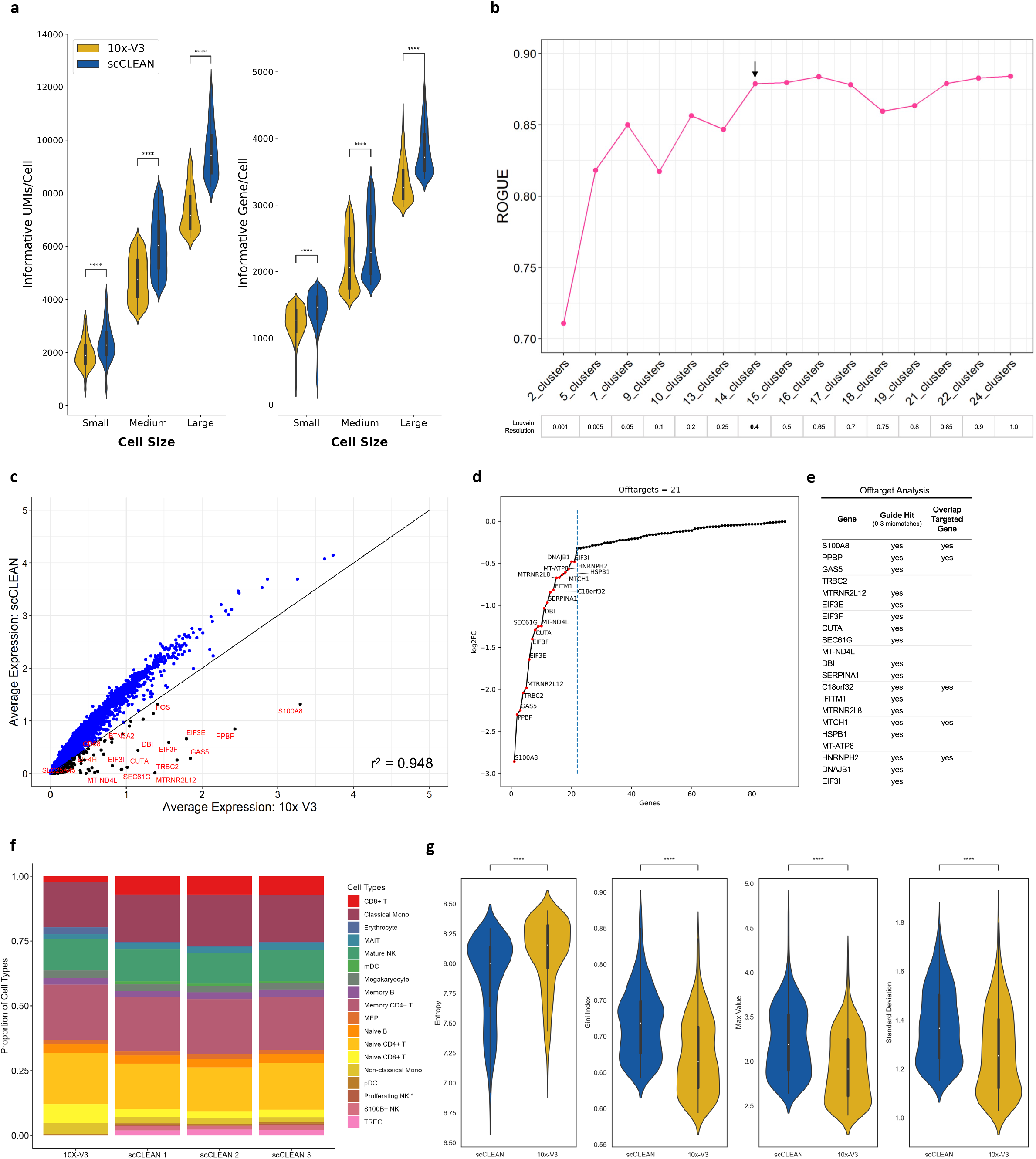
Clustering performance and offtarget analysis. **a**, Comparison of informative UMIs (left) and genes (right) per cell between PBMC samples derived from 10x-V3 (yellow) and scCLEAN (blue) conditions. Due to broad immune cell heterogeneity, cells were separated according to size (3 equivalent sized bins according to metric). **b**, Entropy based ROGUE metric output spanning a range of Louvain resolutions calculated from the standard 10x-V3 sample. The proper clustering resolution (arrow) was selected at the point of saturation and was used to analyze both the 10x-V3 and scCLEAN downstream single cell analysis. **c**, Gene correlation mapping the normalized expression between 10x-V3 and scCLEAN (r^2^ = 0.948). 19,703 genes (88%) increased in expression due to scCLEAN (blue) while 2,569 genes (12%) have higher expression in 10x-V3. **d**, Distinguishing between genes that are a product of random sampling from two libraries of fundamentally different composition and true off-target effects. Genes that don’t follow a linear trend in log2 fold change reduction (negative values represent a decrease in scCLEAN expression relative to 10x-V3) are considered offtarget (n=21) and are automatically detected by a knee bend algorithm. **e**, Quantifying whether the 21 selected genes in this hierarchical selection process are targets of scCLEAN guides or overlap with regions of on-target genes (18/21). **f**, Proportion of identified cell types comparing 10x-V3 to three scCLEAN replicates. 2 replicates identified all 4 additional cell types while 1 replicate only identified 3 out of the 4, omitting proliferating NK cells (labeled with *). **g**, Cell specificity scores (entropy, Gini index, max value, standard deviation) calculated after co-embedding genes onto the latent space within each condition (10x-V3 = yellow, scCLEAN = blue). Top 10% of genes according to each metric were included in analysis and statistical significance was calculated using the Mann-Whitney U rank test with Benjamini-Hochberg correction.

**Supplementary Fig. 2:**
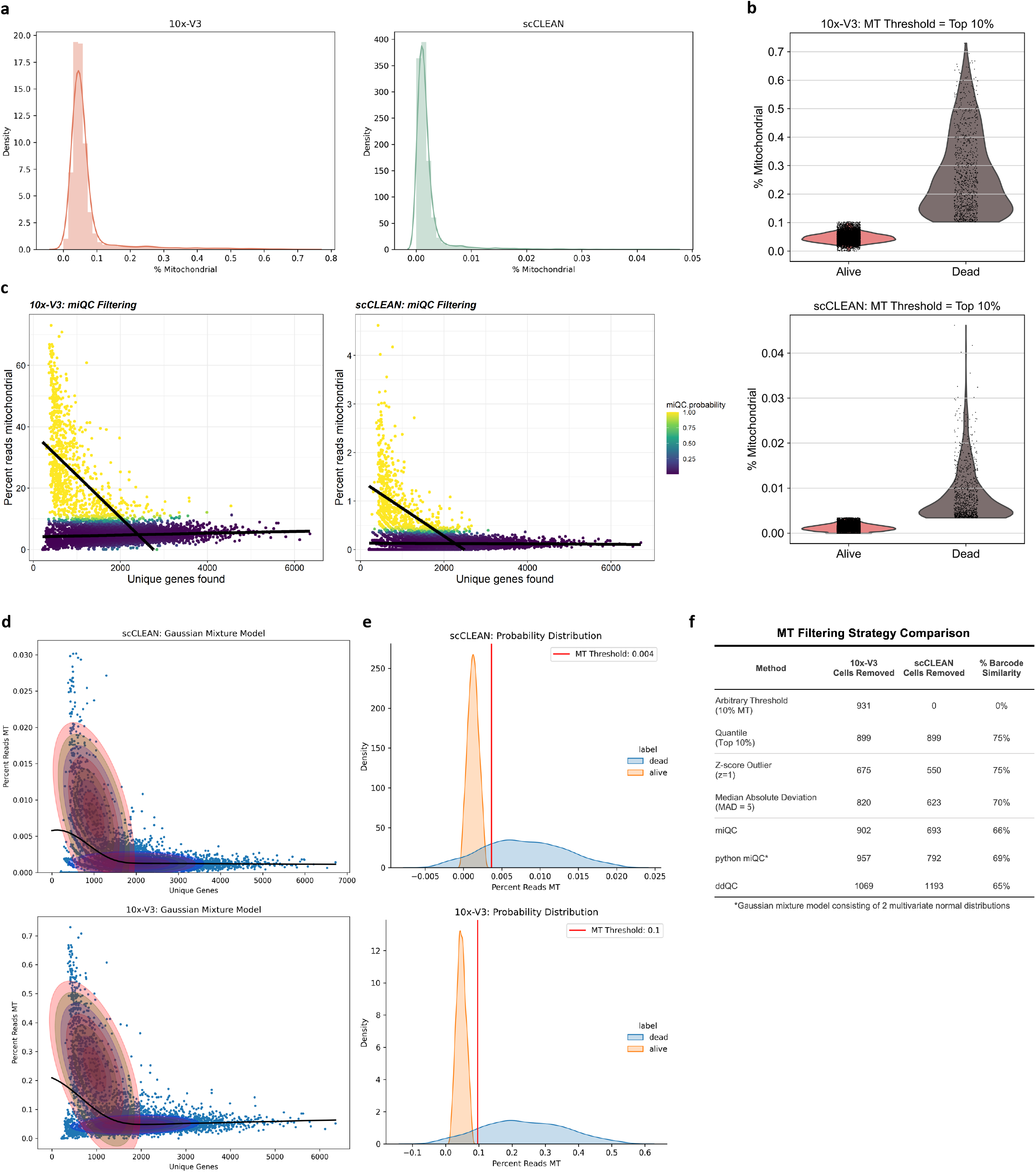
Filtering dead cells using mitochondrial gene expression. **a**, Distribution of cell density according to the percent of mitochondrial (MT) reads comparing a PBMC sample with a 30% ground truth dead cell population (trypan blue staining) prepared with standard 10x-V3 processing (left) and scCLEAN depletion (right). **b**, Violin plots representing the population of dead and alive cells in each condition (10x-V3 = top, scCLEAN = bottom) by setting a threshold removing the top 10% of cells according to MT percentage. **c**, Results of filtering dead cells by using miQC, which is a mixed linear regression model plotting % MT as a function of total number of unique genes. Yellow represents cells classified as dead and subsequently removed. **d**, Represents the same principle as **c**, except models the 2 cell populations as a gaussian mixture. The population with low gene count and high % MT are classified as dead. **e**, Each multivariate normal distribution is plotted (alive = orange, blue = dead) with a 2-sigma confidence region. A proper MT threshold was established by calculating 5 median absolute deviations above the median live cell distribution. **f**, Efficacy of various dead cell filtering techniques. scCLEAN and 10x-V3 samples represent the same cell population so % barcode similarity was calculated to quantify the degree to which the same cells were removed in both conditions according to each filtering technique (scCLEAN filtered barcodes/10x-V3 filtered barcodes).

**Supplementary Fig. 3:**
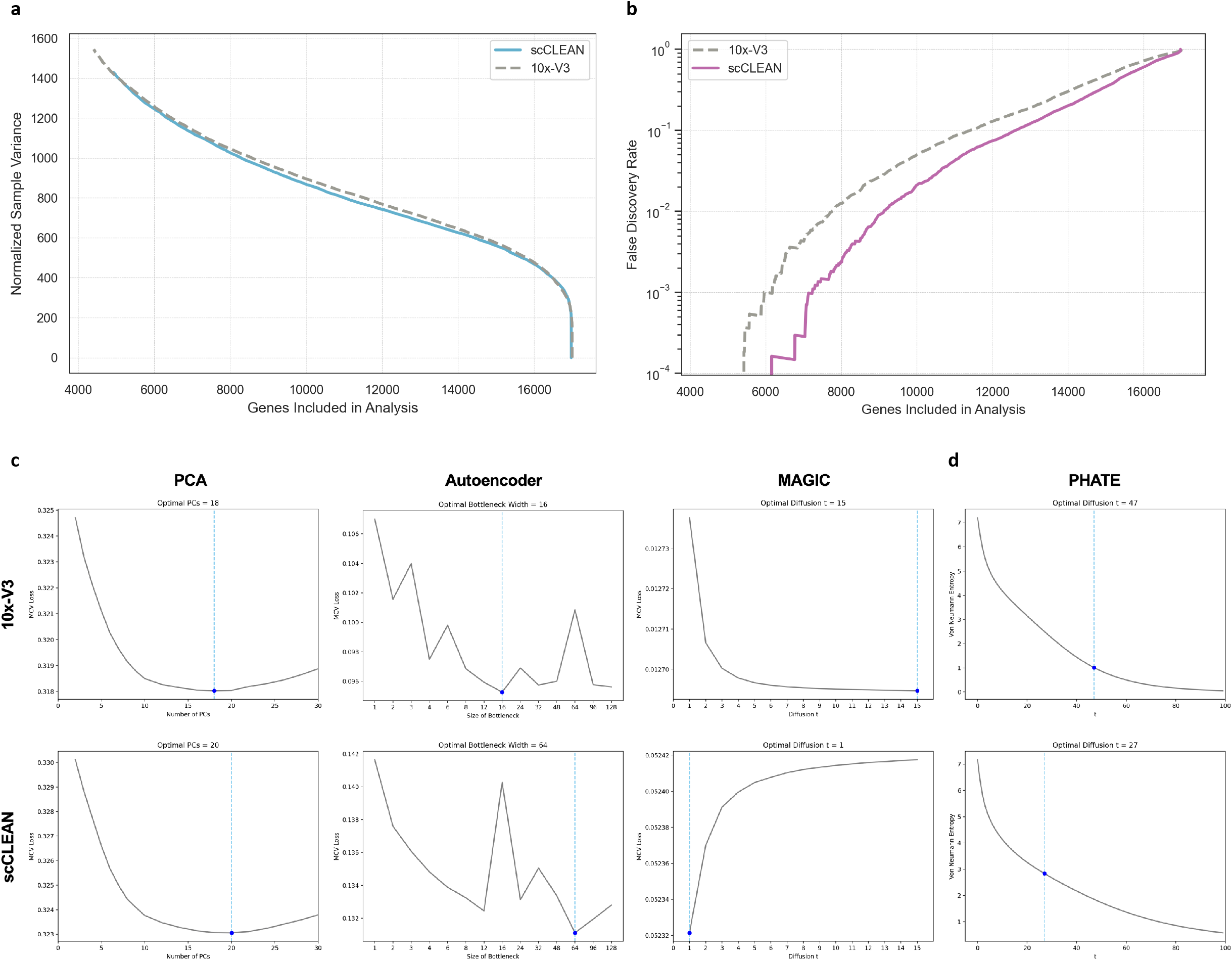
Increased signal to noise enhances dimensionality reduction. **a**, Random matrix theory (RMT) to calculate the change in normalized sample variance as more genes are included in secondary analysis (10x-V3 = gray, scCLEAN = blue). **b**, RMT to plot the false discovery rate as a function of gene inclusion (10x-V3 = gray, scCLEAN = purple). **c**, Comparing the optimized dimensionality reduction (DR) across various techniques (columns) between 10x-V3 (top row) and scCLEAN (bottom row). The optimal key parameter for each technique was determined using molecular cross validation (MCV loss). Left, standard principal component analysis demonstrates increased variance incorporated through DR performance by increasing the number of principal components from 18 (10x-V3) to 20 (scCLEAN). Middle, optimized size of the bottleneck layer for a denoising autoencoder for single cell count data. At the optimal bottleneck width, scCLEAN (64) incorporates more signal due to a wider layer than 10x-V3 (16). Right, proper diffusion parameter t for the standard diffusion-based DR algorithm (MAGIC) illustrates a significantly decreased optimized value from 15 (10x-V3) to 1 (scCLEAN). The larger the value, the greater the genetic heterogeneity is smoothed over in an attempt to reduce noise. **d**, DR performance comparison using a different diffusion-based algorithm (PHATE) and a separate optimization algorithm based on Von Neuman Entropy. The diffusion component is reduced from 47 (10x-V3) to 27 (scCLEAN).

**Supplementary Fig. 4:**
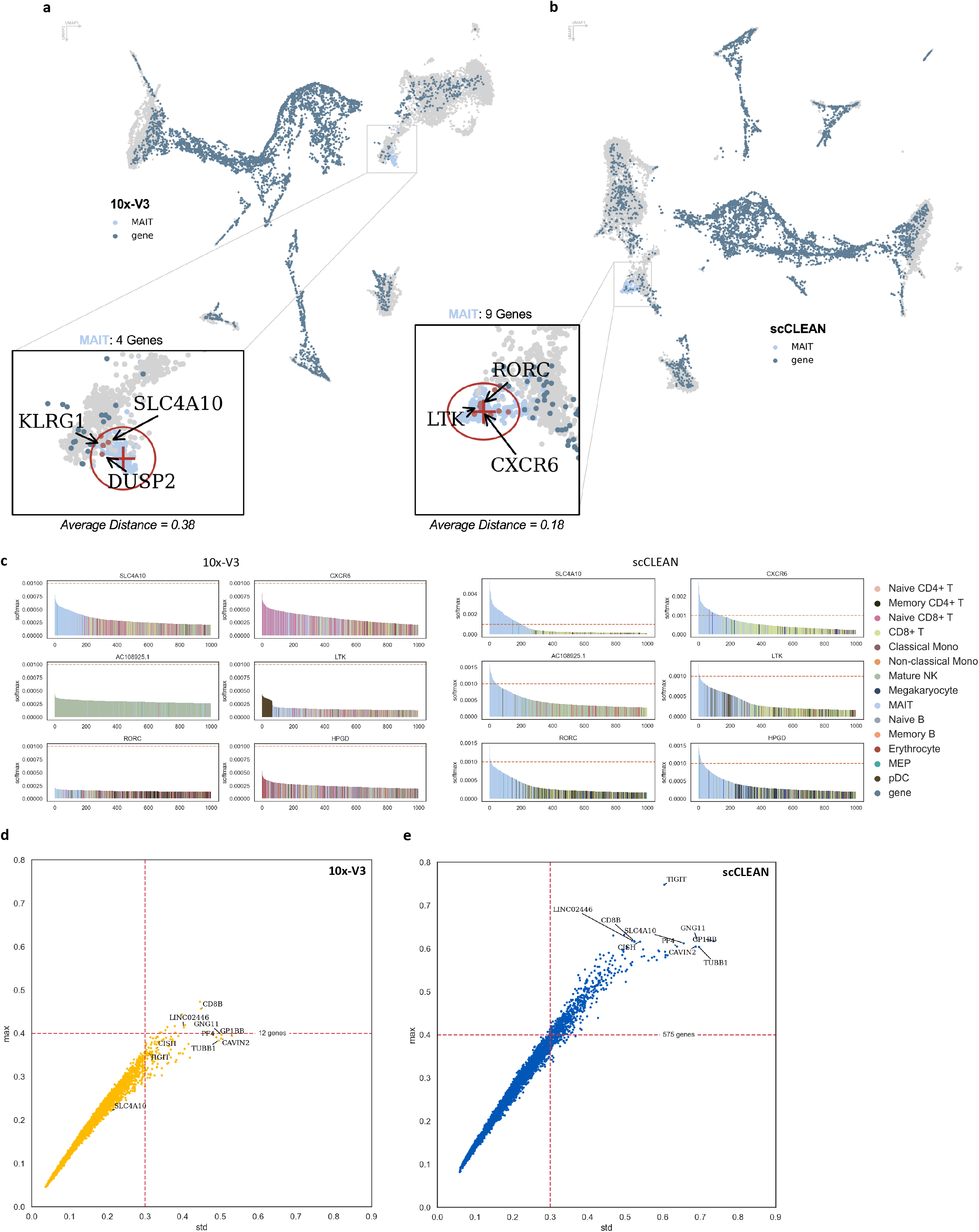
Enhanced specificity of a shared cell type. **a**, Co-embedding of genes onto the 10x-V3 UMAP projection, illustrating cell-type markers for MAIT cells. Center of query automatically selected as point of highest cluster density with a search radius of 0.5 (4 genes identified with an average distance to center of 0.38). **b**, Results of same analysis as **a** but on scCLEAN dataset, identifying 9 genes associated with MAIT cell population with an average distance to center of 0.18 in coordinate space. **c**, Top 6 cell markers identified in scCLEAN for MAIT cells. Plotting the barcodes (colored by cell type, MAIT = light blue) most closely associated (softmax transformation) with each gene. Red dotted line indicates default threshold for gene to cell type specificity. None of the marker genes passed the specificity threshold with 10x-V3. **d**, All 10x-V3 detected genes (yellow) plotted according to MAIT specificity (standard deviation and max value). 12 genes pass a cutoff threshold (std = 0.3 and max=0.4). **e**, For the same cutoff threshold, 575 genes (48-fold increase) are identified as MAIT specific within scCLEAN (blue).

**Supplementary Fig. 5:**
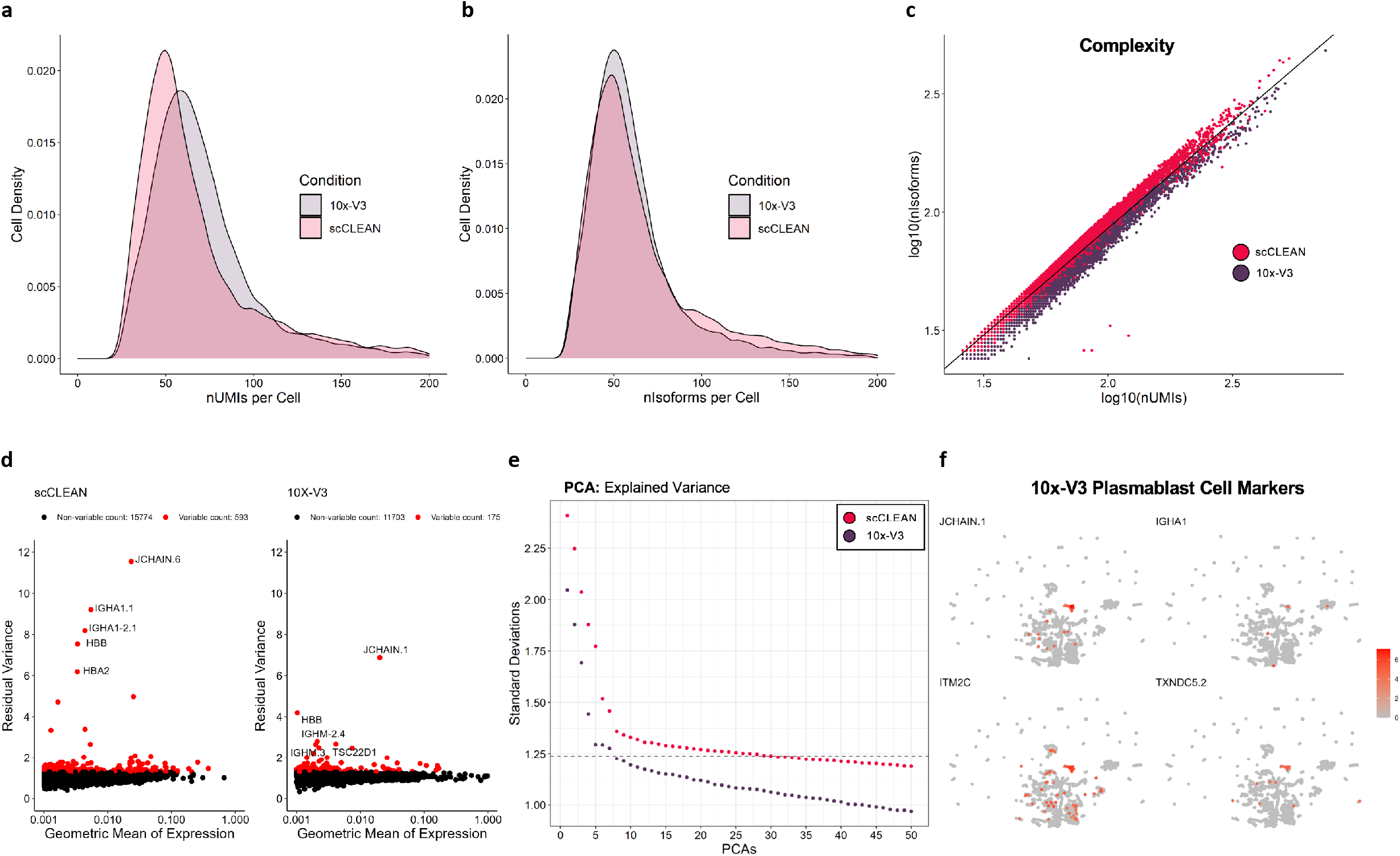
Single cell Iso-seq comparison. **a-f**, Generated from data with isoforms associated with the 255 targeted genes retained. **a**, PBMC iso-seq distribution of the number of UMIs detected per cell (10x-V3 = grey, scCLEAN = pink). **b**, same as **a** except cell density according to number of unique isoforms per cell. **c**, Scatter plot of cell complexity, unique isoforms as a function of UMIs, comparing 10x-V3 (purple) to scCLEAN (red). **d**, Highly variable gene selection identifying 593 genes with scCLEAN (left) and 175 genes with 10x-V3 (right) (3.4-fold increase). **e**, scCLEAN (red) captures higher biological variance with every principal component in comparison to 10x-V3 (purple). Dashed line indicates the number of scCLEAN principal components incorporated into downstream clustering. **f**, 10x-V3 feature plots (UMAP) depicting 4 Plasmablast cell markers.

**Supplementary Fig. 6:**
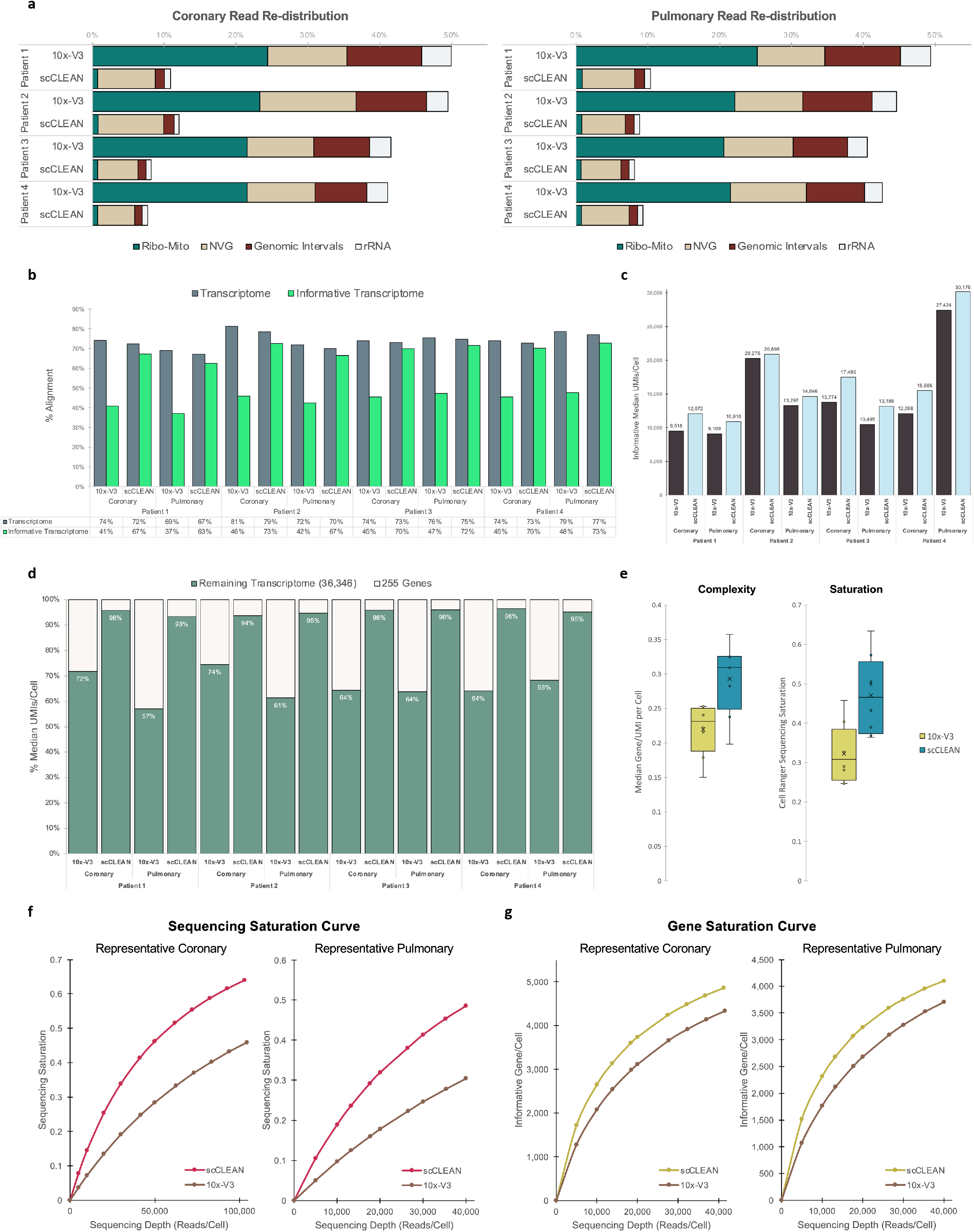
scCLEAN performance evaluation on cohort of vascular smooth muscle cells. **a**, Read re-distribution of 4 targeted regions for removal (Ribo-Mito, NVG, Genomic Intervals, rRNA) within coronary samples (left) and pulmonary samples (right) across 4 patients. **b**, For all samples, alignment rates to the full transcriptome (dark grey) or the informative transcriptome (lime green) corresponding to the exclusion of the 255 targeted un-informative genes. **c**, Median informative UMIs detected per cell across all patient samples comparing 10x-V3 (black) to scCLEAN (light blue). **d**, Ratio of all UMIs per cell corresponding with the 255 targeted genes (tan) and the remaining transcriptome (36,346 genes) (green). **e**, Box and whisker plot illustrating the boost in complexity and sequencing saturation with scCLEAN (blue) relative to 10x-V3 (yellow). **f**, Sequencing saturation as a function of sequencing depth comparing scCLEAN (red) and 10x-V3 (brown) illustrating representative coronary (left) and pulmonary (right) samples. **g**, Saturation curves depicting informative gene detection across sequencing depth between scCLEAN (yellow) and 10x-V3 (brown) illustrating representative coronary (left) and pulmonary (right) samples.

**Supplementary Fig. 7:**
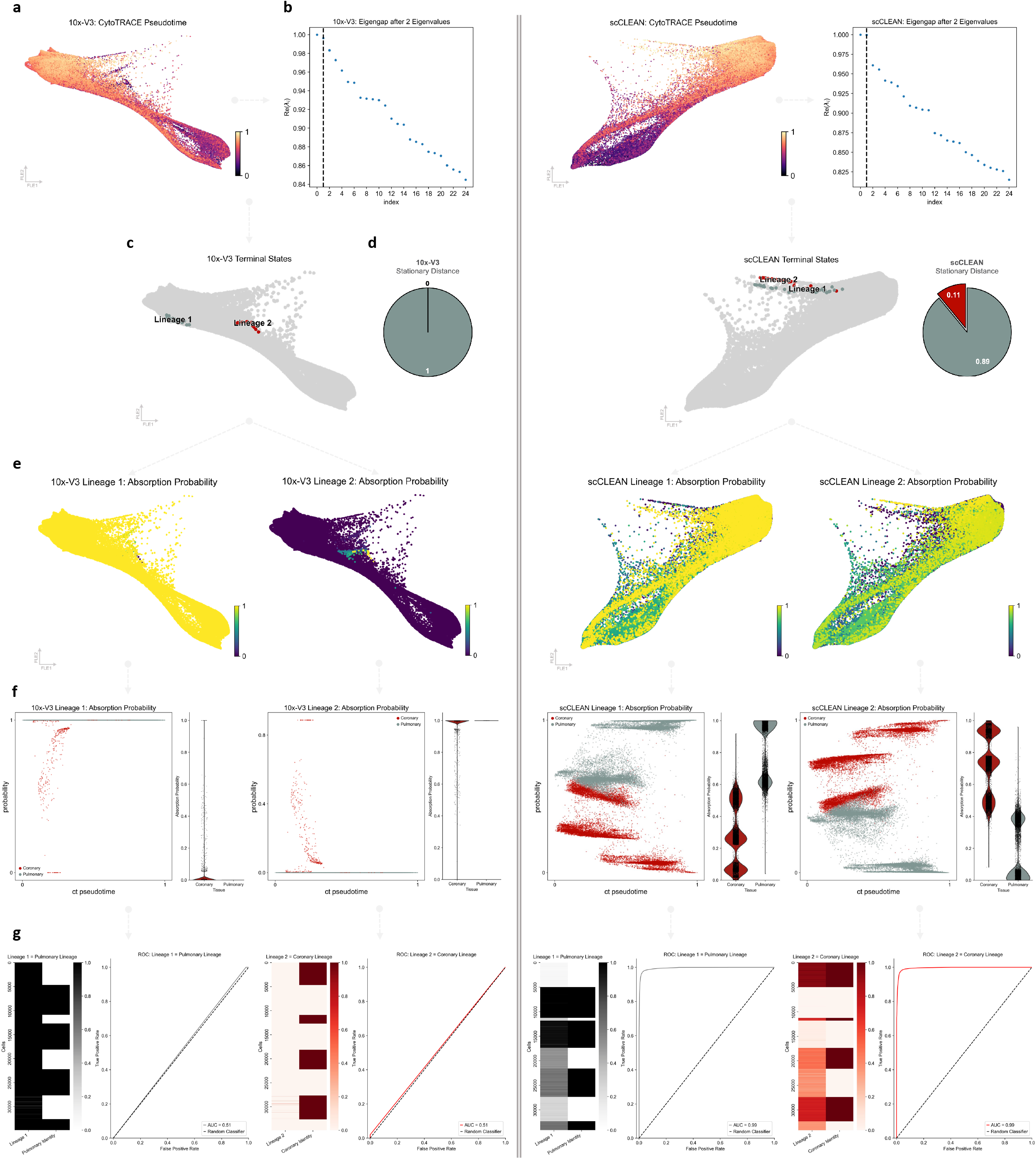
Orthogonal validation of cardiovascular trajectory analysis. **a-g,** Trajectory analysis workflow depicting 10x-V3 (left of line divide) and scCLEAN (right of line divide). **a**, Pseudotime projected on force directed layout plot (FLE), calculated using all genes detected and depicting the transition from early cells (black) to terminal cells (yellow). **b**, Schur decomposition plotting the real components of the top 25 eigenvalues. A gap was calculated after 2 values motivating the calculation of 2 terminal states. **c**, Location of 2 terminal states on FLE projection. **d**, Stationary distance from coarse-grained Markov transition matrix associated with each lineage. For the second lineage identified in both conditions, 10x-V3 cannot identify any transition distance (0) while scCLEAN identifies a relative distance of 0.11. **e**, Probability of each cell belonging to each lineage (absorption probability). Yellow illustrates 100% probability of cell-lineage association while dark blue represents 0%. Greater than 99% of cells within 10x-V3 (left) belong to lineage 1, while in scCLEAN (right), 64% of cells correspond to lineage 1 and 36% of cells correspond to lineage 2 (absorption probability > 0.5). **f**, Each cells lineage absorption probability (lineage 1 = left, lineage 2 = right) plotted as a function of each cells position along the differentiation trajectory (ct pseudotime). Coloring reflects whether the cell was derived from a coronary or pulmonary artery. **g**, First, heat matrix illustrating the probability of each cell belonging to each lineage (lineage 1 = left, lineage 2 =right) paired side by side with the identity of that cell belonging to each tissue. Second, receiver operating characteristic (ROC) depicting the classification performance of identifying each tissue to each lineage. Both lineages of 10x-V3 (left of line divide) have an AUC of 0.51 while the corresponding lineages identified with scCLEAN (right of line divide) have an AUC of 0.99.

## References

1. Shekhar, K. et al. Comprehensive Classification of Retinal Bipolar Neurons by Single-Cell Transcriptomics. Cell 166, 1308–1323.e30 (2016).

2. Trapnell, C. et al. Pseudo-temporal ordering of individual cells reveals dynamics and regulators of cell fate decisions. Nat Biotechnol 32, 381 (2014).

3. Satija, R., Farrell, J. A., Gennert, D., Schier, A. F. & Regev, A. Spatial reconstruction of single-cell gene expression. Nat Biotechnol 33, 495 (2015).

4. Ocone, A., Haghverdi, L., Mueller, N. S. & Theis, F. J. Reconstructing gene regulatory dynamics from high-dimensional single-cell snapshot data. Bioinformatics 31, i89–i96 (2015).

5. Almet, A. A., Cang, Z., Jin, S. & Nie, Q. The landscape of cell–cell communication through single-cell transcriptomics. Curr Opin Syst Biol 26, 12–23 (2021).

6. Armingol, E., Officer, A., Harismendy, O. & Lewis, N. E. Deciphering cell–cell interactions and communication from gene expression. Nat Rev Genet 22, 71–88 (2020).

7. Regev, A. et al. The Human Cell Atlas. Elife 6, (2017).

8. Jones, R. C. et al. The Tabula Sapiens: A multiple-organ, single-cell transcriptomic atlas of humans. Science (1979) 376, (2022).

9. Suo, C. et al. IMMUNOLOGY Mapping the developing human immune system across organs. Science (1979) 376, (2022).

10. Haghverdi, L., Lun, A. T. L., Morgan, M. D. & Marioni, J. C. Batch effects in single-cell RNA sequencing data are corrected by matching mutual nearest neighbours. Nat Biotechnol 36, 421 (2018).

11. Lähnemann, D. et al. Eleven grand challenges in single-cell data science. Genome Biol 21, 1–35 (2020).

12. Hicks, S. C., Townes, F. W., Teng, M. & Irizarry, R. A. Missing data and technical variability in single-cell RNA-sequencing experiments. Biostatistics 19, 562–578 (2018).

13. Phipson, B., Zappia, L. & Oshlack, A. Gene length and detection bias in single cell RNA sequencing protocols. F1000Res 6, (2017).

14. Finak, G. et al. MAST: a flexible statistical framework for assessing transcriptional changes and characterizing heterogeneity in single-cell RNA sequencing data. Genome Biol 16, (2015).

15. Gong, W., Kwak, I. Y., Pota, P., Koyano-Nakagawa, N. & Garry, D. J. DrImpute: Imputing dropout events in single cell RNA sequencing data. BMC Bioinformatics 19, 1–10 (2018).

16. Mou, T., Deng, W., Gu, F., Pawitan, Y. & Vu, T. N. Reproducibility of Methods to Detect Differentially Expressed Genes from Single-Cell RNA Sequencing. Front Genet 10, (2020).

17. Kharchenko, P. v, Silberstein, L. & Scadden, D. T. Bayesian approach to single-cell differential expression analysis. Nat Methods 11, 740–742 (2014).

18. Grün, D., Kester, L. & van Oudenaarden, A. Validation of noise models for single-cell transcriptomics. Nat Methods 11, 637–640 (2014).

19. Hashimshony, T. et al. CEL-Seq2: Sensitive highly-multiplexed single-cell RNA-Seq. Genome Biol 17, 1–7 (2016).

20. Klein, A. M. et al. Droplet barcoding for single-cell transcriptomics applied to embryonic stem cells. Cell 161, 1187–1201 (2015).

21. Zheng, G. X. Y. et al. Massively parallel digital transcriptional profiling of single cells. Nat Commun 8, (2017).

22. Macosko, E. Z. et al. Highly Parallel Genome-wide Expression Profiling of Individual Cells Using Nanoliter Droplets. Cell 161, 1202–1214 (2015).

23. Zhang, L. & Zhang, S. Imputing single-cell RNA-seq data by considering cell heterogeneity and prior expression of dropouts. J Mol Cell Biol 13, 29–40 (2021).

24. Carolina LeoteID, A., WuID, X. & BeyerID, A. Regulatory network-based imputation of dropouts in single-cell RNA sequencing data. PLoS Comput Biol 18, (2022).

25. Qiu, P. Embracing the dropouts in single-cell RNA-seq analysis. Nat Commun 11, (2020).

26. Saliba, A. E., Westermann, A. J., Gorski, S. A. & Vogel, J. Single-cell RNA-seq: advances and future challenges. Nucleic Acids Res 42, 8845–8860 (2014).

27. Kolodziejczyk, A. A., Kim, J. K., Svensson, V., Marioni, J. C. & Teichmann, S. A. The technology and biology of single-cell RNA sequencing. Mol Cell 58, 610–620 (2015).

28. van Dijk, E. L., Jaszczyszyn, Y. & Thermes, C. Library preparation methods for nextgeneration sequencing: Tone down the bias. Exp Cell Res 322, 12–20 (2014).

29. Lahens, N. F. et al. IVT-seq reveals extreme bias in RNA sequencing. Genome Biol 15, (2014).

30. Mamanova, L. et al. FRT-seq: amplification-free, strand-specific transcriptome sequencing. Nat Methods 7, 130–132 (2010).

31. Khan, S. & Kaihara, K. A. Single-cell RNA-sequencing of peripheral blood mononuclear cells with ddSEQ. Methods in Molecular Biology 1979, 155–176 (2019).

32. Wenger, A. M. et al. Accurate circular consensus long-read sequencing improves variant detection and assembly of a human genome. Nat Biotechnol 37, 1155–1162 (2019).

33. Volden, R. et al. Improving nanopore read accuracy with the R2C2 method enables the sequencing of highly multiplexed full-length single-cell cDNA. Proceedings of the National Academy of Sciences 115, 9726–9731 (2018).

34. Chakraborty, R. et al. Targeting smooth muscle cell phenotypic switching in vascular disease. JVS Vasc Sci 2, 79–94 (2021).

35. Svensson, V., da Veiga Beltrame, E. & Pachter, L. A curated database reveals trends in single-cell transcriptomics. Database 2020, (2020).

36. Ilicic, T. et al. Classification of low quality cells from single-cell RNA-seq data. Genome Biol 17, 1–15 (2016).

37. Luecken, M. D. & Theis, F. J. Current best practices in single-cell RNA-seq analysis: a tutorial. Mol Syst Biol 15, e8746 (2019).

38. Linderman, G. C. Dimensionality Reduction of Single-Cell RNA-Seq Data. Methods in Molecular Biology 2284, 331–342 (2021).

39. Johnson, E. M., Kath, W. & Mani, M. EMBEDR: Distinguishing signal from noise in single-cell omics data. Patterns 3, 100443 (2022).

40. Pinder, C. L. et al. Isolation and Characterisation of Antigen-Specific Plasmablasts Using a Novel Flow Cytometry-Based Immunoglobulin Capture Assay (ICA). Journal of Immunology 199, 4180 (2017).

41. Hu, Z. et al. Single-Cell Transcriptomic Atlas of Different Human Cardiac Arteries Identifies Cell Types Associated With Vascular Physiology. Arterioscler Thromb Vasc Biol 41, 1408–1427 (2021).

42. Miano, J. M., Fisher, E. A. & Majesky, M. W. Fate and State of Vascular Smooth Muscle Cells in Atherosclerosis. Circulation 143, 2110–2116 (2021).

43. Pedroza, A. J. et al. Single-Cell Transcriptomic Profiling of Vascular Smooth Muscle Cell Phenotype Modulation in Marfan Syndrome Aortic Aneurysm. Arterioscler Thromb Vasc Biol 40, 2195–2211 (2020).

44. Sorokin, V. et al. Role of Vascular Smooth Muscle Cell Plasticity and Interactions in Vessel Wall Inflammation. Front Immunol 11, 3053 (2020).

45. Yap, C., Mieremet, A., de Vries, C. J. M., Micha, D. & de Waard, V. Six Shades of Vascular Smooth Muscle Cells Illuminated by KLF4 (Krüppel-Like Factor 4). Arterioscler Thromb Vasc Biol 41, 2693–2707 (2021).

46. Pokhilko, A. et al. Targeted single-cell RNA sequencing of transcription factors enhances the identification of cell types and trajectories. Genome Res 31, 1069–1081 (2021).

47. Singh, M. et al. High-throughput targeted long-read single cell sequencing reveals the clonal and transcriptional landscape of lymphocytes. Nat Commun 10, (2019).

48. Replogle, J. M. et al. Combinatorial single-cell CRISPR screens by direct guide RNA capture and targeted sequencing. Nat Biotechnol 38, 954–961 (2020).

49. Schraivogel, D. et al. Targeted Perturb-seq enables genome-scale genetic screens in single cells. Nat Methods 17, 629 (2020).

50. Dynerman, D. et al. Designing and implementing programmable depletion in sequencing libraries with DASHit. bioRxiv (2020) doi:10.1101/2020.01.12.891176.

51. Loi, D. S. C., Yu, L. & Wu, A. R. Effective ribosomal RNA depletion for single-cell total RNA-seq by scDASH. PeerJ 9, (2021).

52. Hardigan, A. A. et al. CRISPR/Cas9-targeted removal of unwanted sequences from small-RNA sequencing libraries. Nucleic Acids Res 47, (2019).

53. Gu, W. et al. Depletion of Abundant Sequences by Hybridization (DASH): Using Cas9 to remove unwanted high-abundance species in sequencing libraries and molecular counting applications. Genome Biol 17, 1–13 (2016).

54. Bennett, M. R., Sinha, S. & Owens, G. K. Vascular smooth muscle cells in atherosclerosis. Circ Res 118, 692 (2016).

55. Soehnlein, O. & Libby, P. Targeting inflammation in atherosclerosis — from experimental insights to the clinic. Nature Reviews Drug Discovery 2021 20:8 20, 589–610 (2021).

56. Ezan, J. et al. FrzA/sFRP-1, a secreted antagonist of the Wnt-Frizzled pathway, controls vascular cell proliferation in vitro and in vivo. Cardiovasc Res 63, 731–738 (2004).

57. Wang, S. H. et al. Pigment epithelium-derived factor reduces the PDGF-induced migration and proliferation of human aortic smooth muscle cells through PPARγ activation. Int J Biochem Cell Biol 44, 280–289 (2012).

58. Ji, Y. et al. Pharmacological Targeting of Plasminogen Activator Inhibitor-1 Decreases Vascular Smooth Muscle Cell Migration and Neointima Formation. Arterioscler Thromb Vasc Biol 36, 2167–2175 (2016).

59. Chou, E. L., Cardenas, C. L., Conrad, M. F. & Lindsay, M. E. Vascular Smooth Muscle Cells Release Protease Nexin-1 in Cell Models of Atherosclerosis and Human Carotid Tissue. J Vasc Surg 72, e262–e263 (2020).

60. Ashino, T., Yamamoto, M., Yoshida, T. & Numazawa, S. Redox-sensitive transcription factor Nrf2 regulates vascular smooth muscle cell migration and neointimal hyperplasia. Arterioscler Thromb Vasc Biol 33, 760–768 (2013).

61. He, X., Cheng, R., Benyajati, S. & Ma, J. X. PEDF and its roles in physiological and pathological conditions: implication in diabetic and hypoxia-induced angiogenic diseases. Clin Sci 128, 805 (2015).

62. Guimaraes, J. C. & Zavolan, M. Patterns of ribosomal protein expression specify normal and malignant human cells. Genome Biol 17, 1–13 (2016).

63. Chen, A. et al. Spatiotemporal transcriptomic atlas of mouse organogenesis using DNA nanoball-patterned arrays. Cell 185, 1777–1792.e21 (2022).

64. Cho, C. S. et al. Microscopic examination of spatial transcriptome using Seq-Scope. Cell 184, 3559–3572.e22 (2021).

65. Rodriques, S. G. et al. Slide-seq: A scalable technology for measuring genome-wide expression at high spatial resolution. Science (1979) 363, 1463–1467 (2019).

66. Vickovic, S. et al. High-definition spatial transcriptomics for in situ tissue profiling. Nat Methods 16, 987 (2019).

67. Ståhl, P. L. et al. Visualization and analysis of gene expression in tissue sections by spatial transcriptomics. Science 353, 78–82 (2016).

68. Lee, J., Hyeon, D. Y. & Hwang, D. Single-cell multiomics: technologies and data analysis methods. Experimental & Molecular Medicine 2020 52:9 52, 1428–1442 (2020).

69. Robinson, J. T. et al. Integrative Genomics Viewer. Nat Biotechnol 29, 24 (2011).

70. Li, B. et al. Cumulus provides cloud-based data analysis for large-scale single-cell and single-nucleus RNA-seq. Nat Methods 17, 793 (2020).

71. Franzén, O., Gan, L. M. & Björkegren, J. L. M. PanglaoDB: a web server for exploration of mouse and human single-cell RNA sequencing data. Database 2019, 46 (2019).

72. Hafemeister, C. & Satija, R. Normalization and variance stabilization of single-cell RNA-seq data using regularized negative binomial regression. Genome Biol 20, 1–15 (2019).

73. Becht, E. et al. Dimensionality reduction for visualizing single-cell data using UMAP. Nat Biotechnol 37, 38–47 (2018).

74. Germain, P.-L., Lun, A., Meixide, C. G., Macnair, W. & Robinson, M. D. Doublet identification in single-cell sequencing data using scDblFinder. F1000Res 10, 979 (2021).

75. Liu, B. et al. An entropy-based metric for assessing the purity of single cell populations. Nature Communications 2020 11:1 11, 1–13 (2020).

76. Jin, S. et al. Inference and analysis of cell-cell communication using CellChat. Nature Communications 2021 12:1 12, 1–20 (2021).

77. Hie, B., Cho, H., DeMeo, B., Bryson, B. & Berger, B. Geometric Sketching Compactly Summarizes the Single-Cell Transcriptomic Landscape. Cell Syst 8, 483–493.e7 (2019).

78. Ekiz, H. A., Conley, C. J., Stephens, W. Z. & O’Connell, R. M. CIPR: a web-based R/shiny app and R package to annotate cell clusters in single cell RNA sequencing experiments. BMC Bioinformatics 21, 191 (2020).

79. Monaco, G. et al. RNA-Seq Signatures Normalized by mRNA Abundance Allow Absolute Deconvolution of Human Immune Cell Types. Cell Rep 26, 1627 (2019).

80. Aparicio, L., Bordyuh, M., Blumberg, A. J. & Rabadan, R. A Random Matrix Theory Approach to Denoise Single-Cell Data. Patterns 1, 100035 (2020).

81. Li, X. et al. Deep learning enables accurate clustering with batch effect removal in singlecell RNA-seq analysis. Nature Communications 2020 11:1 11, 1–14 (2020).

82. Chen, H., Ryu, J., Vinyard, M. E., Lerer, A. & Pinello, L. SIMBA: SIngle-cell eMBedding Along with features. bioRxiv (2022) doi:10.1101/2021.10.17.464750.

83. O’Leary, N. A. et al. Reference sequence (RefSeq) database at NCBI: current status, taxonomic expansion, and functional annotation. Nucleic Acids Res 44, D733–D745 (2016).

84. Korsunsky, I. et al. Fast, sensitive and accurate integration of single-cell data with Harmony. Nature Methods 16, 1289–1296 (2019).

85. Lange, M. et al. CellRank for directed single-cell fate mapping. Nature Methods 19, 159–170 (2022).

86. Gulati, G. S. et al. Single-cell transcriptional diversity is a hallmark of developmental potential. Science (1979) 367, 405–411 (2020).

87. Reuter, B., Weber, M., Fackeldey, K., Röblitz, S. & Garcia, M. E. Generalized Markov State Modeling Method for Nonequilibrium Biomolecular Dynamics: Exemplified on Amyloid β Conformational Dynamics Driven by an Oscillating Electric Field. J Chem Theory Comput 14, 3579–3594 (2018).

88. Faure, L., Soldatov, R., Kharchenko, P. v & Adameyko, I. scFates: a scalable python package for advanced pseudotime and bifurcation analysis from single cell data. bioRxiv (2022) doi:10.1101/2022.07.09.498657.

